# Machine Learning Ensemble Directed Engineering of Genetically Encoded Fluorescent Calcium Indicators

**DOI:** 10.1101/2023.04.13.536801

**Authors:** Sarah J. Wait, Michael Rappleye, Justin Daho Lee, Netta Smith, Andre Berndt

**Author notes:** Conflicts of interest.

## Abstract

Real-time monitoring of biological activity can be achieved through the use of genetically encoded fluorescent indicators (GEFIs). GEFIs are protein-based sensing tools whose biophysical characteristics can be engineered to meet experimental needs. However, GEFIs are inherently complex proteins with multiple dynamic states, rendering optimization one of the most challenging problems in protein engineering. Most GEFIs are engineered through trial-and-error mutagenesis, which is time and resource-intensive and often relies on empirical knowledge for each GEFI. We applied an alternative approach using machine learning to efficiently predict the outcomes of sensor mutagenesis by analyzing established libraries that link sensor sequences to functions. Using the GCaMP calcium indicator as a scaffold, we developed an ensemble of three regression models trained on experimentally derived GCaMP mutation libraries. We used the trained ensemble to perform an in silico functional screen on a library of 1423 novel, untested GCaMP variants. The mutations were predicted to significantly alter the fluorescent response, and off-rate kinetics were advanced for verification in vitro. We found that the ensemble’s predictions of novel variants’ biophysical characteristics closely replicated what we observed of the variants in vitro. As a result, we identified the novel ensemble-derived GCaMP (eGCaMP) variants, eGCaMP and eGCaMP+, that achieve both faster kinetics and larger fluorescent responses upon stimulation than previously published fast variants. Furthermore, we identified a combinatorial mutation with extraordinary dynamic range, eGCaMP2+, that outperforms the tested 6th, 7th, and 8th generation GCaMPs. These findings demonstrate the value of machine learning as a tool to facilitate the efficient prescreening of mutants for functional characteristics. By leveraging the learning capabilities of our ensemble, we were able to accelerate the identification of promising mutations and reduce the experimental burden associated with screening an entire library. Machine learning tools such as this have the potential to complement emerging high-throughput screening methodologies that generate massive datasets, which can be tedious to analyze manually. Overall, these findings have significant implications for developing new GEFIs and other protein-based tools, demonstrating the power of machine learning as an asset in protein engineering.

## Introduction

Genetically encoded fluorescent indicators (GEFIs) are protein-based sensors that allosterically link fluorescent proteins to protein domains that bind with specific ligands. Changes in fluorescence intensity upon ligand binding can be quantified spatiotemporally, allowing researchers to monitor ligands such as intracellular second messengers or neuromodulators in freely moving animals^1^. Today, GEFIs are essential tools in neuroscience, with sensors already developed for calcium, dopamine, norepinephrine, endocannabinoids, and opioids, amongst others^2–11^. To align each sensor’s biophysical properties to experimental needs, GEFIs require extensive engineering (i.e., mutation). However, most current engineering techniques, such as trial-and-error mutagenesis, are time and resource intensive. We require novel approaches that ameliorate the experimental burden associated with GEFI engineering. Here, we use data-driven interrogation of a mutational library to investigate the sequence-function relationships of GEFIs and predict the functional characteristics of novel mutants.

Machine learning (ML) algorithms are valuable tools in protein engineering, demonstrating their ability to understand complex interactions without prior knowledge of intricate structure-function relationships.^12^. These algorithms have been employed to engineer enzymes, fluorescent proteins, and optogenetic tools with varying levels of ML-pipeline complexity. ^13–18^. In this study, we integrate the strengths of these previous examples into a novel and enhanced approach. Firstly, our approach is based on the sequence-function relationship, ensuring that the resulting model can be applied to any mutation library. This versatility allows for broader adaptability across various protein engineering problems. Secondly, we utilize multiple models in parallel, implementing an ensembling process that informs our final predictions with increased accuracy and robustness. This methodology offers an advantage, as it remains applicable regardless of the availability of a molecular structure. By combining these elements, we developed an improved ML-based approach with the potential to broadly impact protein engineering. The resulting pipeline is not only adaptable and versatile but also offers better predictive capabilities, paving the way for more efficient engineering of proteins.

We selected the calcium indicator GCaMP as a protein sensor scaffold to develop this platform. GCaMP is a chimeric protein that consists of circularly-permuted GFP (cpGFP) fused to calmodulin (CaM) and calmodulin-binding peptide (CBP). GCaMP sensors have been widely adopted and have seen several improvements to optimize their capabilities^2–6, 19, 20^. As a result, data from the *in vitro* characterizations of >1000 mutants became publicly available^4, 5^. Using this data, we developed a stacked ensemble capable of predicting *in vitro* functional characteristics of previously untested GCaMP mutants. As a result, we identified mutations that accelerate the off-rate kinetics and increase the fluorescent responses of jGCaMP7s. Our study demonstrates that ML ensembles can effectively learn from complex mutational datasets and that we can harness their predictive power to prescreen mutation libraries for enhancing biophysical properties. This quality is increasingly important to complement the growing suite of high-throughput protein engineering methodologies, streamlining the analysis process and further advancing the field.

## Results

### Description of Variant Library, Computational Approach, and Predictions on Novel Sequences

We used public data from two previous publications to form our GCaMP variant libraries, which consisted of 1078 characterized mutants and their *in vitro* functional characteristics derived from cultured neuron screening^4, 5^. Within the variant library, we focused on the fluorescent response (ΔF/F_0_) to one action potential (AP) stimulation (1AP ΔF/F_0_) and decay kinetics of the fluorescent response (τ_1/2,_ decay half-time after 10 APs) (***Figure 1A***). When normalized to GCaMP6s as the baseline for the target attributes (i.e., 1AP ΔF/F_0_ and τ_1/2_ = 1.0), we can see a broad distribution of variant capabilities and mutation locations within the GCaMP structure (**Figure 1B, C; Supp. Fig. 1A**). We found that the sequence similarity is not deterministic for either the fluorescent or kinetic response, as seen by the variability in mutation performance regardless of GCaMP generation (**Supp. Fig. 1B, D**).

**Figure 1:**
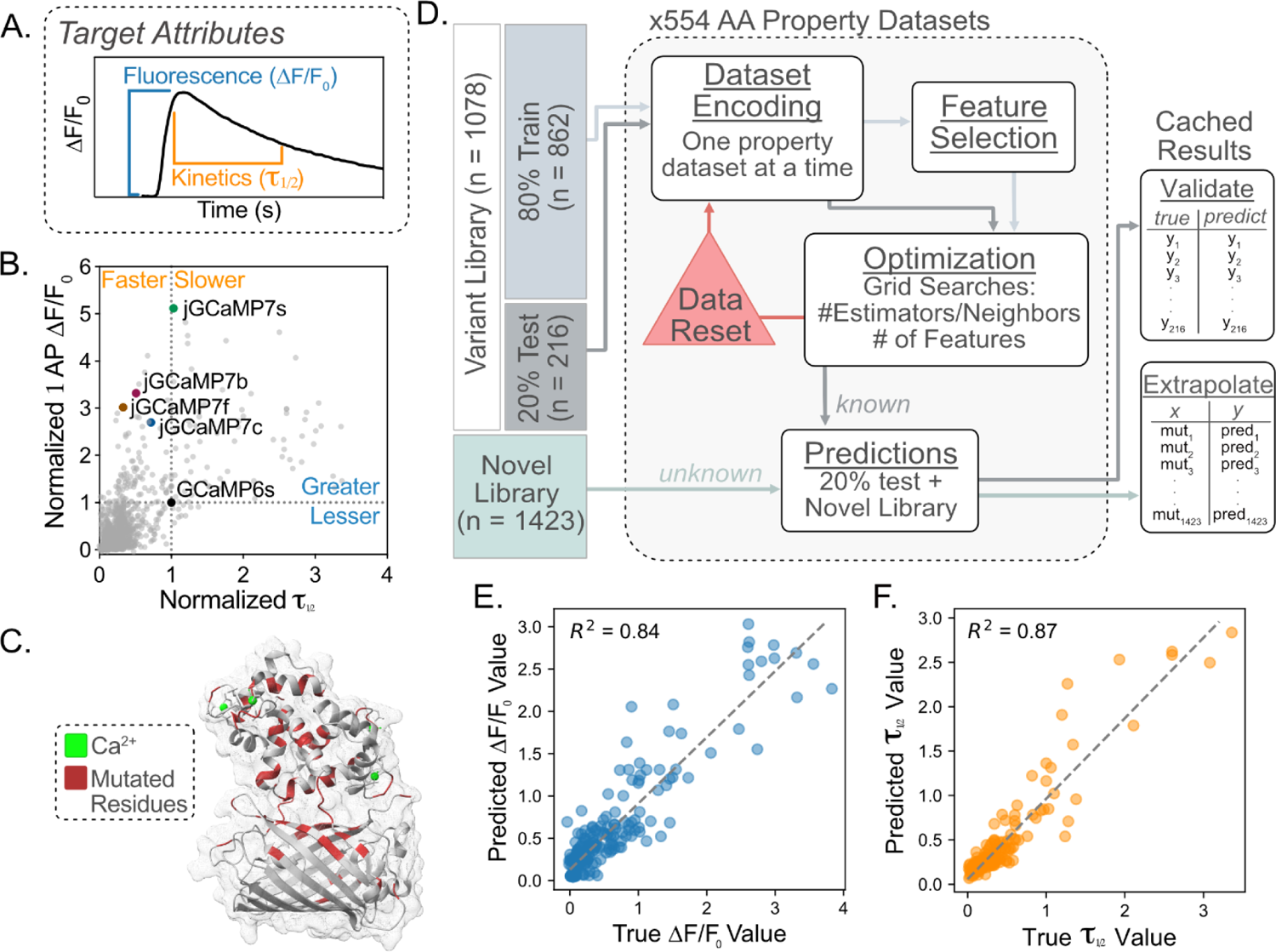
Description of Variant Library, Computational Approach, and Ensemble Cross-Validation. **A.** Description of the biophysical attributes of the GCaMP sensor targeted for engineering. Fluorescence (ΔF/F_0_) is the change between the baseline and maximal fluorescence upon calcium sensing. Kinetics (τ_1/2_) refers to the decay from maximum ΔF/F_0_ to half-maximal ΔF/F_0_. **B.** Scatter plot depicts the 1AP ΔF/F_0_ by the τ_1/2_ for each of the 1078 variants in the variant library^4, 5^. Each value was normalized to GCaMP6s as 1.0 for 1AP ΔF/F_0_ and τ_1/2_. Published variants are indicated with colored dots and text labels. **C.** Crystal structure of GCaMP3-D380Y (RCSB: 3SG3, gray) with 75 residues (red) in which mutation information exists in the variant library^4, 5^. These 75 residues indicate the positions used to form the novel library. **D.** Overview of model training schema. The variant library^4, 5^ was split randomly into an 80% training set and a 20% testing set. The data was encoded using the AAINDEX property datasets. The train set underwent feature selection before being optimized using a grid search of key hyperparameters for each model. The optimized model was used to form predictions on the 20% test set and the novel library. The final predictions for both the test set and novel library were cached for downstream analysis. **E.** Cross-validation of the fluorescence ensemble. The scatter plot x-axis represents the true ΔF/F_0_ value for each variant in the test set, and the y-axis represents the predictions made by the ensemble of the variants in the test set. The dotted line depicts perfect agreement between true values and predicted values. R^2^ value denotes the goodness of fit of scatter data with the dotted line. **F.** Cross-validation of the kinetic ensemble. The scatter plot’s x-axis represents the true τ_1/2_ value contained for each variant in the test set and the y-axis represents the predictions made by the ensemble of the variants in the test set. The dotted line depicts perfect agreement between true values and predicted values. R^2^ value denotes the goodness of fit of scatter data with the dotted line.

Before model training, the variants in the library were randomly assigned to training and testing sets at an 80/20 ratio for downstream cross-validation, where the mean values between the train and test sets were not significantly different in either the fluorescence or kinetics library (**Supp. Fig. 1C, E**). To improve prediction capabilities, we performed a stacked ensemble comprising a random forest regressor (RFR), K-neighbors regressor (KNR), and multi-layer perceptron network regressor (MPNR)^21, 22^. We encoded each position in the GCaMP sequence in the train and test sets using values that quantify amino acid (AA) properties such as size, polarity, hydrophobicity, etc. (554 property datasets x 3 models). (**Figure 1D**). The cross-validation R^2^ value was used to benchmark each AA property’s ability to predict the functional ability of the withheld test set (**Supp. Figure 2A**). Only the top five AA property datasets for each model were considered in the final ensemble to reduce the computational burden, time, and storage space. Interestingly, the underlying AA properties that led to high R^2^ values were associated with hydrophobicity for the fluorescence library and conformation for the kinetics library (**Supp. Fig. 2B, C, D; Supp. Table 1, 2**).

**Figure 2:**
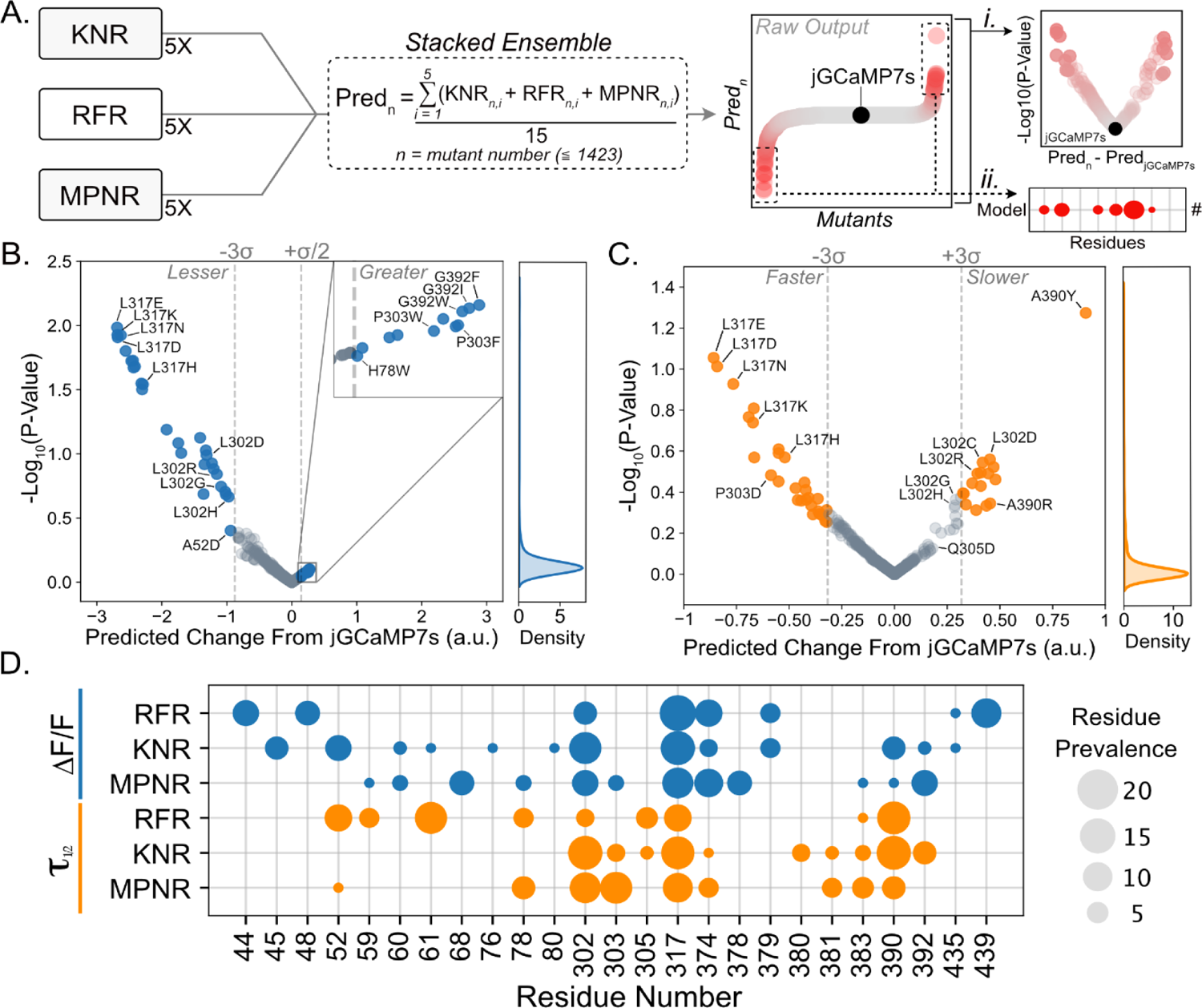
Predictions Derived From the Ensembles Led to Mutations of Interest for In Vitro Verification. **A.** Brief description of prediction analysis. From each model, the predictions from the top five property datasets were combined in the stacked ensemble. The stacked ensemble predictions were formed by averaging the predictions from the 15 contributor models for each variant (Pred_n_) in the novel library. The raw output is thus the prediction (Pred_n_) for each mutant, with a prediction for jGCaMP7s as a benchmark. The volcano plots were formed by subtracting the benchmark jGCaMP7s prediction from the variant prediction (x-axis) and P-values were derived by performing an unpaired t-test between the 15 predictions for variant_n_ and the 15 predictions for jGCaMP7s (**i**). The bubble plot indicates the prevalence (or the number of occurrences) of each residue that resides in the top 2.5% and bottom 2.5% of predictions (**ii**). **B.** Volcano plots depicting the ensemble’s prediction for a given mutation change in fluorescent response from jGCaMP7s (x-axis) and the log_10_(P-value) of the given prediction. P-values were derived by performing an unpaired t-test on ensemble prediction (15 models) for jGCaMP7s and given mutation. Kernel density estimation (right) depicts the spread of log_10_(P-values) obtained. **C.** Volcano plots depicting the ensemble’s prediction for given mutations change kinetic capability from jGCaMP7s (x-axis) and the log_10_(P-value) of the given prediction. P-values were derived using an unpaired t-test on ensemble prediction (15 models) for jGCaMP7s and given mutation. Kernel density estimation (right) depicts the spread of log_10_(P-values) obtained. **D.** Bubble plot depicting the number of times each residue (x-axis) appeared in the top 2.5% and bottom 2.5% of predicted values for each regressor that comprise each ensemble.

The ensemble’s predictions for each mutation are the average response from the 15 models (5 AA properties x 3 models). During cross-validation, the ensembles for fluorescence and kinetics achieved R^2^ values greater than 0.80 for predictions made on the test dataset (**Figure 1E, F**). The fluorescence ensemble achieved a higher R^2^ value than any of the models contributing to the prediction, pointing to the beneficial collaborative effect of ensembling (**Supp. Fig. 2C**). Importantly, we found that the addition of the amino acid property information improves the model’s ability to generalize the variant library, as evidenced by label encoded libraries achieving R^2^ values of 0.66/0.68 and one-hot encoded libraries achieving R^2^ values of 0.70/0.66 for the fluorescence (1AP ΔF/F_0_) and kinetics (τ_1/2_) predictions, respectively (**Supp. Fig. 2C**).

### Identification of Mutations of Interest From Ensemble Predictions

We utilized the trained ensembles to predict a novel library’s fluorescence and kinetics capabilities. This library was created by taking jGCaMP7s and substituting each of the 75 positions previously mutated in the variant library with the remaining 19 amino acids (**Figure 1C**). After removing redundant variants, the novel library contained 1423 untested variants. We calculated the ‘Predicted Change From jGCaMP7s’ by subtracting the average predicted value of jGCaMP7s from the predicted value for each mutant. We performed an unpaired t-test between the 15 predictions made for each mutant (one from each contributor model) and the 15 predictions made for jGCaMP7s within the same library. These two metrics allowed us to isolate mutations whose predicted value differs significantly from jGCaMP7s (**Figure 2Ai**). Next, we isolated the residues in each library whose mutations had the strongest positive or negative impact on fluorescence and kinetics (**Figure 2Aii**). From these normalized value predictions, we can ascertain how mutations were predicted to affect the biophysical characteristics of jGCaMP7s in both the fluorescence and kinetics (**Figure 2B, C**). In our model training, the jGCaMP7s sequence was purposely withheld. Nevertheless, the ensemble prediction ranked the base jGCaMP7s sequence within the top 15% of variants for high fluorescence readout. Consequently, the ensemble predicted most variants to have a decreased fluorescent response compared to jGCaMP7s. Variants such as L317E, L317K, L317N, L317D, and L317H were all predicted to have a decreased fluorescent response (<-2.2 a.u.) compared to jGCaMP7s, while variants such as G392F, G392I, and G392W were all predicted to have an increased (>0.25 a.u.) fluorescent response (**Figure 2B**). In the kinetics library, L317E, L317D, L317N, and L317K were all predicted to decay faster (<-0.6 a.u.) than jGCaMP7s, while variants such as A390Y, L302D, and L302C were all predicted to decay slower (>0.3 a.u.) than jGCaMP7s (**Figure 2C**). The variants discussed above all fell outside 99.7% (±3σ) of −log_10_(P-Values), except for high fluorescence predictions, indicating that the 15 contributing models displayed confidence in the effect of the mutation (±3σ, fluorescence: 0.612, kinetics: 0.242) (**Figure 2B, C**).

We found that 22% and 18% of the impactful residues in the fluorescence and kinetics libraries, respectively, were L317 predictions (**Figure 2D**), despite only 1.3% of variants in the novel library harboring an L317 mutation. Similarly, L302 predictions accounted for 14% and 16% of the impactful residues of the fluorescence and kinetics libraries, respectively (**Figure 2D**). L317 is located on the interface between CaM and CBP, while L302 is part of the linker between CaM and cpGFP-both are key positions within GCaMP (**Supp. Fig. 3A, B, C**). In contrast, residue A390 was found to be 4.5 times more impactful in the kinetics predictions than in the fluorescence predictions. Like L317, A390 is located on the interface between CaM and CBP but on the opposing side (***Supp. Fig. 3D***). Impactful residues for each biophysical property also tended to cluster. For instance, the kinetics library displays 38% prediction prevalence surrounding residue clusters Y380, R381, R383, and L302, P303, Q305. The prevalence of these residues is 2.38x higher in kinetics predictions than the fluorescence predictions. These residues are located close to each other in 3D space representing the residue linker and one of the inward loops of CaM (**Supp. Fig. 3E**). Within the fluorescence predictions, residue clusters N44, K45, H48, V52, and M374, M378, K379 displayed 31% prediction prevalence, 3.9x higher than the kinetics library. Interestingly, when mapping residues H48, V52, L317, M374, M378, and K379 back onto the crystal structure, we observed that all of these residues face inward toward one another, suggesting that they may be involved in interactions essential for the fluorescent response (**Supp. Fig. 3F**). These observations allow us to concentrate mutation efforts on key residues and identify specific residues or residue interactions that may be most advantageous to target for each biophysical characteristic.

**Figure 3:**
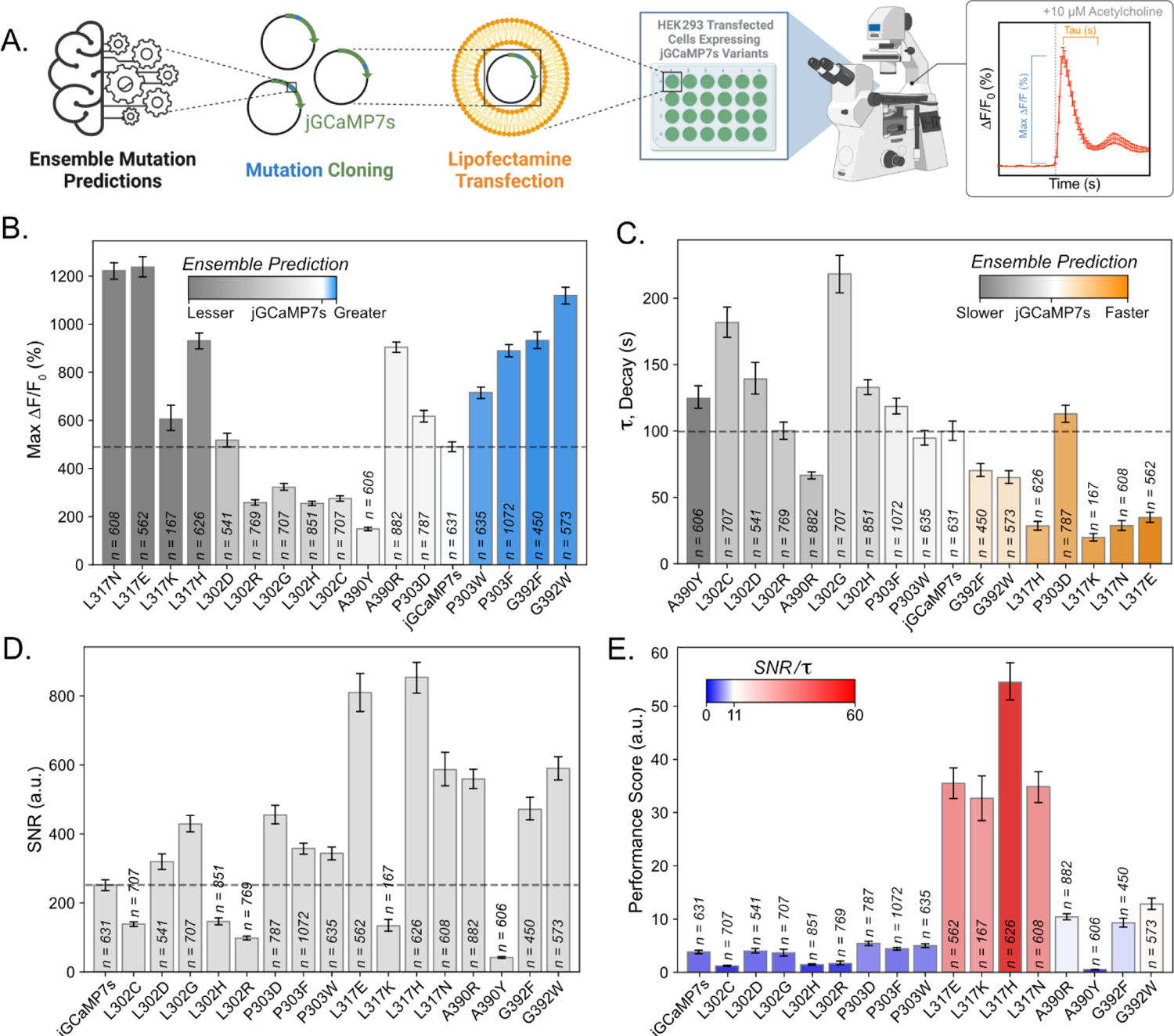
Gq/IP3 Assay in HEK293 Cells To Validate Ensemble Predictions. **A.** Brief description of the methods contained in the figure. Mutation predictions isolated from the ensemble are used as the basis for downstream variant analysis. Variants of interest are cloned into the jGCaMP7s (Addgene, #104463) backbone. These variants are then transfected into HEK293 cells using lipofectamine transfection. Forty-eight hours post-transfection, cells are time-course imaged using an epifluorescent microscope. The stimulation protocol contains a period to collect baseline fluorescence, a bath addition of 10 µM acetylcholine, and a decay period. Visual representations of the qualifications in **B./C.** are found on the representative response trace. **B.** Max fluorescent responses (*Eq.1*) that were obtained from each mutant of jGCaMP7s expressed in HEK293 cells and stimulated with ten µM acetylcholine. Heat mapping demonstrates the ensemble’s prediction of the given mutation’s performance *in vitro*. Mutations are sorted in order of the ensemble’s predicted performance. (n = number of cells quantified; bars depict mean + bootstrapped 95% ci^42^) **C.** Decay values (τ, tau, *Eq.4*) obtained from each mutant of jGCaMP7s expressed in HEK293 cells and stimulated with ten µM acetylcholine. Heat mapping demonstrates the ensemble’s prediction of the given mutation’s performance *in vitro*. Mutations are sorted in order of the ensemble’s predicted performance. (n = number of cells quantified; bars depict mean + bootstrapped 95% ci^42^). **D.** Signal-to-noise ratio (SNR, *Eq.2*) of each mutant of jGCaMP7s expressed in HEK293 cells and stimulated with ten µM acetylcholine. (n = number of cells quantified; bars depict mean + bootstrapped 95% ci^42^). **E.** Performance score, consisting of the SNR/τ (*Eq.2/Eq.4*), obtained from each mutant of jGCaMP7s expressed in HEK293 cells and stimulated with ten µM acetylcholine. Heat mapping highlights the highest-scoring mutants or those with high ΔF/F_0_ (%) responses and fast decay speeds. (n = the number of cells quantified; bars depict mean + 95% bootstrapped ci^42^).

### In Vitro Performance of Ensemble Predictions

We tested mutations predicted by the ML-ensemble to enhance biophysical properties by stimulating HEK293 cells with acetylcholine^2, 3, 23^. This process activates calcium channels in the endoplasmic reticulum (ER) through G_q_/IP_3_ coupled pathways^24, 25^ (**Figure 3A**). Mutations predicted to have a greater ΔF/F_0_ than jGCaMP7s (P303F, P303W, G392F, and G392W) all achieved >130% increase in fluorescence over jGCaMP7s (**Figure 3B, Supp. Table 3A**). We also found three variants (L302G, L302H, and L302R) that satisfied their predicted decrease in fluorescent response, with an average of 1.75x lower fluorescent response (**Figure 3B, Supp. Table 3A**). However, we found that the L317 mutants, which were predicted to have a decreased fluorescent response displayed the opposite characteristic *in vitro*. For example, all four L317 mutants achieved 2x greater ΔF/F_0_ than jGCaMP7s. A retrospective analysis of the previous GCaMP mutations showed that variants containing 317N, 317E, 317K, or 317H saw almost a complete reduction of the fluorescent response (**Supp. Fig. 4A, B**). Accordingly, we found that the L317H mutation in jGCaMP7f led to the predicted reduction of fluorescent response (**Supp. Fig. 4C**). This reflects findings in the Dana *et al.* 2019 dataset, in which variant 10.1035 (jGCaMP7f L317H) saw a 95% reduction in ΔF/F_0_ to 1AP stimuli compared to 10.9210 (jGCaMP7f)^5^. We speculate that this learned association is why the ensemble predicted mutations at L317 are detrimental to the sensor’s fluorescent response. Regardless, we found multiple examples of mutations that led to the altered fluorescent response we were aiming to tune.

**Figure 4:**
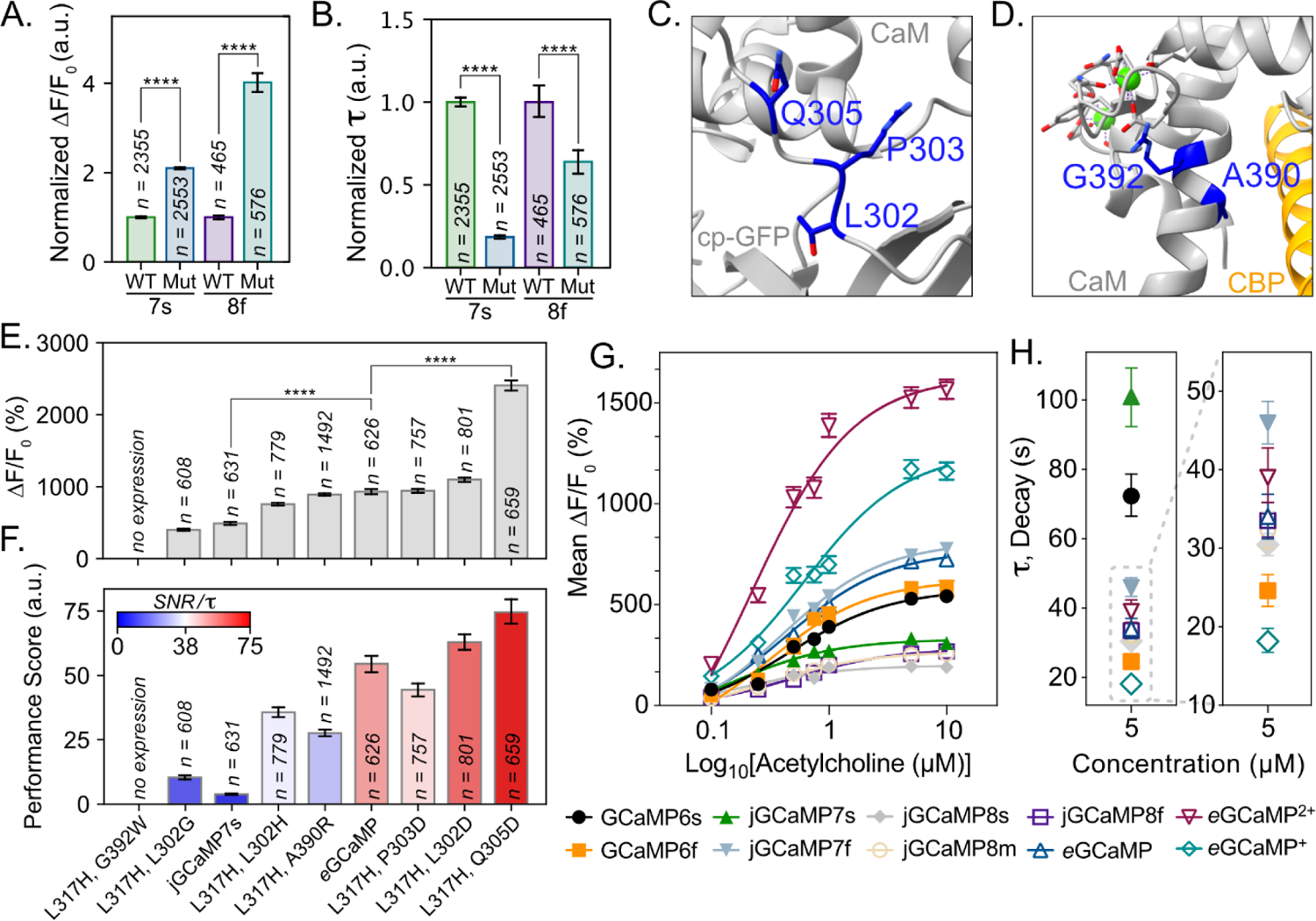
Mutation Transfer and Combinatorial Mutation For The Identification of eGCaMP^+^ and eGCaMP^2+^. **A.** Max fluorescent responses (*Eq.1*) obtained from each variant indicated on the x-axis, expressed in HEK293 cells and stimulated with ten µM acetylcholine. Wild Type (WT) indicates the parent construct of either jGCaMP7s (7s) or jGCaMP8f (8f). Mutation (Mut) indicates the parental construct with the addition of L317H in jGCaMP7s and A289H in jGCaMP8f. Each parental/variant pair is normalized to the base construct mean = 1.0 (n = the number of cells quantified; bars depict mean + bootstrapped 95% ci^42^; **** = <0.0001 (unpaired t-test)). jGCaMP8f A289H is called *e*GCaMP*^+^* in Figure 4G. and **4H. B.** Decay values (τ, tau, *Eq.4*) obtained from each variant indicated on the x-axis, expressed in HEK293 cells and stimulated with ten µM acetylcholine. Wild Type (WT) indicates the parent construct of either jGCaMP7s (7s) or jGCaMP8f (8f). Mutation (Mut) indicates the parental construct with the addition of L317H in jGCaMP7s and A289H in jGCaMP8f. Each parental/variant pair is normalized to the base construct mean = 1.0 (n = the number of cells quantified; bars depict mean + bootstrapped 95% ci^42^; **** = <0.0001 (unpaired t-test)). jGCaMP8f A289H is called *e*GCaMP*^+^* in Figure 4G. and **4H.** Crystal structure of GCaMP3-D380Y (RCSB: 3SG3, gray) with Q305 and linker residues P303 and L302 colored in dark blue with sidechains visible. CaM and cpGFP labels are included to orient linker locations. **C.** Crystal structure of GCaMP3-D380Y (RCSB: 3SG3, gray) with A390 and G392 colored dark blue with sidechains visible. Bound Ca^2+^ (green spheres) in the EF-Hand motifs and the CBP (orange) are included. **D.** Max fluorescent responses (*Eq.1*) obtained from each combinatorial variant of jGCaMP7s expressed in HEK293 cells and stimulated with ten µM acetylcholine. Mutations are sorted in order of ΔF/F_0_ performance and identified on the x-axis of D. (n = the number of cells quantified; bars depict mean + bootstrapped 95% ci^42^; **** = <0.0001 (unpaired t-test)). **E.** Performance score, consisting of the SNR/τ (*Eq.2/Eq.4*), obtained from each combinatorial variant of jGCaMP7s expressed in HEK293 cells and stimulated with 10 µM acetylcholine. Mutations are sorted in order of ΔF/F_0_ performance. (n = number of cells quantified; bars depict mean + bootstrapped 95% ci^42^) jGCaMP7s L317H Q305D is called *e*GCaMP*^2+^* in Figure 4G. and **4H. F.** Fluorescent responses (ΔF/F_0,_ *Eq. 1*) of indicated variant, expressed in HEK293 cells and stimulated with different acetylcholine concentrations (x-axis). Plotted points indicate the mean ΔF/F_0_ response for each variant to indicated stimuli, and error bars display the SEM. The solid line depicts the non-linear fit of scatter data. Additional information on plotted points is included in Supplementary Table 3. **G.** Kinetic decay (τ, tau, *Eq.4*) of indicated variant, expressed in HEK293 cells and stimulated with 5 µM acetylcholine. Plotted points indicate the mean tau for each variant to the indicated stimuli, and error bars display the SEM. Additional information on plotted points is included in Supplementary Table 3.

The mutations that changed kinetics largely aligned with the ensemble predictions (**Figure 3C**). Variants P303D, L317E, L317H, L317K, L317N, G392F, and G392W were predicted to accelerate decay kinetics. Of these variants, 85% showed shorter decay times than jGCaMP7s, with L317K displaying a decay time that was 5x faster than jGCaMP7s (**Figure 3C, Supp. Table 3B**). Additionally, 71% of the variants predicted to decrease decay (L302C, L302D, L302G, L302H, L302R, A390R, A390Y) demonstrated the predicted behavior, with L302G exhibiting a decay time 2.18 times longer than jGCaMP7s (**Figure 3C, Supp. Table 3B**).

For subsequent experiments, we focussed on mutations that increased ΔF/F_0_ and accelerated decay kinetics, as these biophysical characteristics could improve the detection of fast calcium signaling, such as those found in neurons firing APs. We found that the variants with large fluorescent responses, including G392W, G392F, P303F, P303W, L317N, L317K, L317E, and L317H, maintained an average signal-to-noise ratio (SNR, Eq. 2) 1.5x greater than jGCaMP7s (**Figure 3D, Supp. Table 3C**). To highlight variants with large fluorescence and fast kinetics, we created a performance score by dividing SNR by the tau value (*Eq. 2/Eq. 4*) (**Figure 3E**). L317E, L317K, L317H, and L317N achieved performance scores on average 10.28x greater than jGCaMP7s (**Supp. Table 3D**). Among them, L317H had the highest performance score of 54.49 (a.u.), 14.23x greater than jGCaMP7s. Based on this assessment, we selected the jGCaMP7s L317H variant for further characterization and named it “ensemble-GCaMP” (eGCaMP). These in vitro results demonstrate that the ensemble could effectively predict sensor functionality, significantly reducing the experimental burden required to identify variants with desired biophysical properties.

### Combinatorial Mutations and Mutation Transfer Led to the Identification of eGCaMP^+^ and eGCaMP^2+^

We introduced the 317H mutation into jGCaMP8f^6^ to test if the beneficial effects could similarly alter divergent GCaMP iterations. Residue L317 in jGCaMP7s is located in a conserved region of CaM and is equivalent to A289 in jGCaMP8f (**Supp. Fig. 5A**). The A289H mutation on jGCaMP8f improved the fluorescent response 4x over jGCaMP8f (**Figure 4A**). jGCaMP8f A289H also showed 36% faster decay than jGCaMP8f (**Figure 4B**). The fast decay kinetics combined with large fluorescent responses provide a promising variant that we named ensemble GCaMP + (*e*GCaMP^+^), which we advanced for further downstream testing.

**Figure 5:**
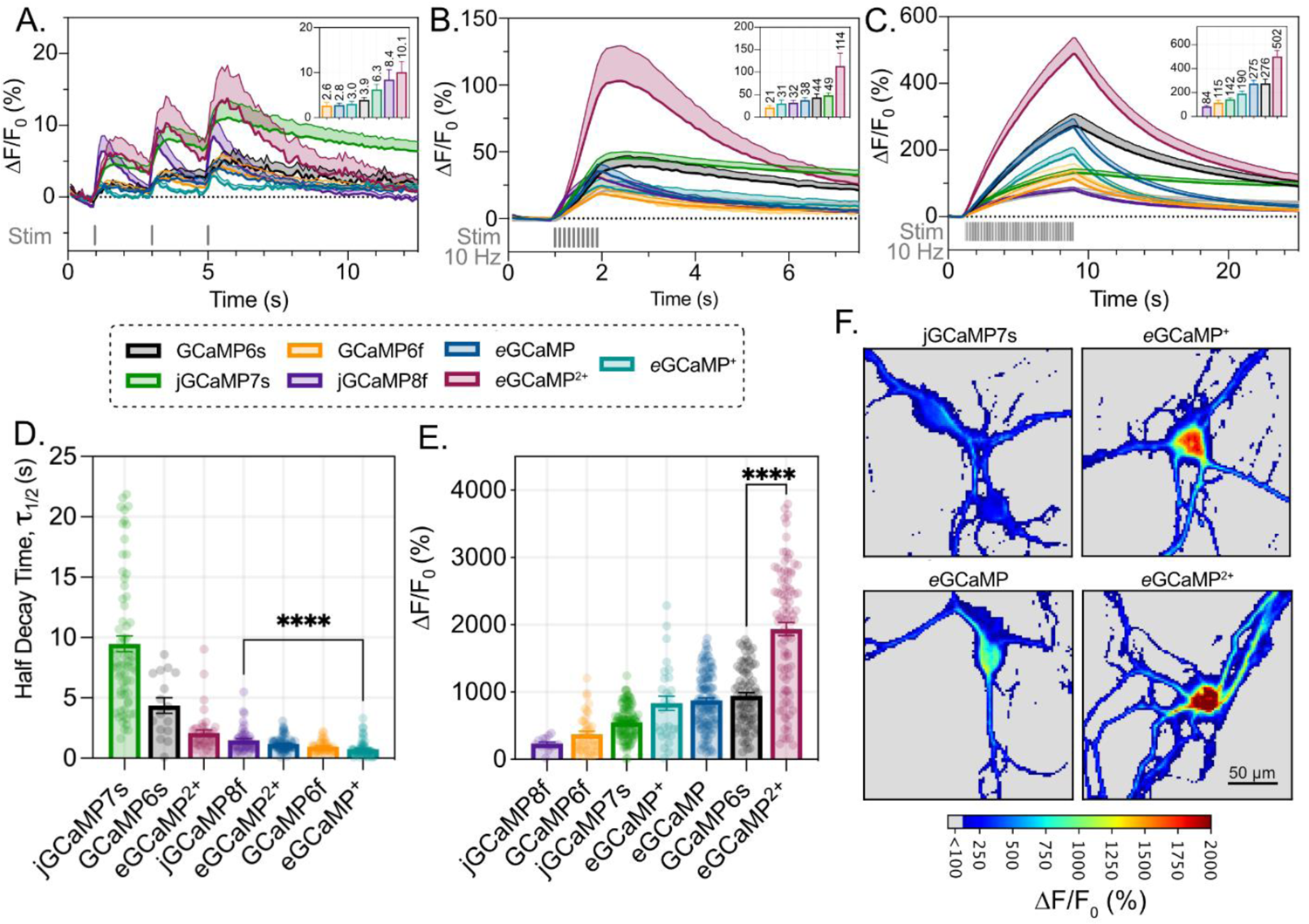
eGCaMP, eGCaMP^+^, and eGCaMP^2+^ Fluorescence and Kinetics Characteristics in Primary Neurons. **A.** ΔF/F_0_ (%) recordings of each variant to 1 AP stimuli applied at 1 Hz over 3 seconds (lines depict mean, shading depicts SEM). The applied stimulus is shown in gray. Graph inset displays max ΔF/F_0_ (%) of each variant to first applied AP (mean + SEM, above-bar annotations = mean). **B.** ΔF/F_0_ (%) recordings of each variant to 10 AP stimuli applied at 10 Hz over 1 second (lines depict mean, shading depicts SEM). The applied stimulus is shown in gray. Graph inset displays max ΔF/F_0_ (%) of each variant to 10 AP stimulus (mean + SEM, above-bar annotations = mean). **C.** ΔF/F_0_ (%) recordings of each variant to 80 AP stimuli applied at 10 Hz over 8 seconds (lines depict mean, shading depicts SEM). The applied stimulus is shown in gray. Graph inset displays max ΔF/F_0_ (%) of each variant to 80 AP stimulus (mean + SEM, above-bar annotations = mean). **D.** Half decay time values after 10 AP stimuli, scatter depicts neurons quantified. (bars depict mean + SEM; * = 0.045 (Unpaired t-test, Two-tailed)). **E.** Maximum ΔF/F_0_ (%) achieved after stimulation with 40 mM KCl. (bars depict mean + SEM; **** = < 0.0001 (Unpaired t-test, Two-tailed)). Representative images of maximal fluorescent response to 40 mM KCl stimulation variant indicated above image. Heat Mapping displays ΔF/F_0_ (%) achieved by each pixel. (Scale bar = 50 µm).

Next, we tested a select combination of additional mutations on *e*GCaMP. For example, we chose the jGCaMP7s variants L302D, P303D, A390R, and G392W for their increased ΔF/F_0_ *in vitro* (**Figure 3B**). Other mutants were selected based on their locations. Namely, L302 and P303 are key functional residues in the linker between cpGFP and CaM^3, 26^(**Figure 4C**). Residue G392 forms a hydrogen bond with residue G398, which lies in one of the EF-hand domains and has been previously observed to influence the Ca^2+^ affinity^3, 27^ (**Supp. Fig. 3D**), and A390 lies on the interaction face between CaM and CBP (**Figure 4D**). We tested Q305 due to its proximity to the linker residues (**Figure 4C**), hydrogen bonding interactions with Y380 (**Supp. Fig. 3E**), and prevalence in the impactful residues for kinetics (**Figure 2D**).

All combinations, except for L317H/G392W, led to functional proteins (**Figure 4E, F; Supp. Fig. 5B, C**).On Average, all variants exhibited decay times 5.0x faster than jGCaMP7s (**Supp. Fig. 5B; Supp. Table 4B**). Within the tested variants, 50% displayed equal or improved fluorescent response to that of *e*GCaMP (**Figure 4E; Supp. Table 4A**). We observed the largest ΔF/F_0_ in the L317H/Q305D mutation, with an almost 2.5-fold increase in ΔF/F_0_ over *e*GCaMP and a 5-fold increase over jGCaMP7s (**Figure 4E; Supp. Table 4A**). The variant also achieved the highest performance score (i.e., large SNR, fast decay) of all variants, a 1.36x fold increase over *e*GCaMP (**Figure 4F; Supp. Fig. 5B, C; Supp. Table 4D**). We chose the jGCaMP7s L317H/Q305D for further characterization and named it ensemble-GCaMP^2+^ (*e*GCaMP^2+^).

We benchmarked the baseline fluorescence, ΔF/F0 capabilities, and kinetic decays of eGCaMP, eGCaMP^2+^, and eGCaMP^+^ against published variants at different acetylcholine concentrations, including widely used constructs such as GCaMP6s, GCaMP6f, jGCaMP7s, jGCaMP7f, jGCaMP8s, jGCaMP8m, and jGCaMP8f^4–6^. Using c-terminally red fluorescent protein (RFP) tagged constructs, we found that *e*GCaMP, *e*GCaMP^+^, and *e*GCaMP^2+^ maintained higher dynamic ranges and SNRs but have lower baseline fluorescences than GCaMP6s, jGCaMP7s, and jGCaMP8f (**Figure 4F; Supp. Figure 6C, D**). In the acetylcholine concentration curve, we found that the three ensemble variants demonstrated impressive fluorescent responses compared to the previously published constructs (**Figure 4G**). At every tested concentration, *e*GCaMP^+^ and *e*GCaMP^2+^ maintained larger ΔF/F_0_s than all previously published variants (**Figure 4G; Supp. Table 5**). For example, eGCaMP^2+^ achieved 2.5 times greater ΔF/F0s at 0.1µM acetylcholine than the highest performing published variant, with decay times comparable to jGCaMP7f (**Figure 4G, H; Supp. Table 5**). Additionally, the decay time of *e*GCaMP^+^ was the fastest of all tested variants (46% faster than jGCaMP8f, its parental construct) while the maximum ΔF/F_0_, was second only to *e*GCaMP^2+^ (**Figure 4G, H; Supp. Table 5**). The *e*GCaMP achieved a ΔF/F_0_ close to jGCaMP7f but with a 26% faster decay (**Figure 4G, H; Supp. Table 5**).

### eGCaMP, eGCaMP^+^, and eGCaMP^2+^ Performance in Primary Neurons

Next, we tested eGCaMP variants in cultured primary rat cortical neurons while stimulating using extracellular electrical fields^4, 5, 28^. *e*GCaMP^2+^ displayed a ΔF/F_0_ of 10.1% in response to 1 AP stimuli, similar to amplitudes obtained by jGCaMP8f (**Figure 5A; Supp. Table 6A**). *e*GCaMP^2+^’s impressive response amplitudes became more apparent with increasing numbers of elicited action potentials. At 10 AP, *e*GCaMP^2+^ achieved 2.34x greater response than jGCaMP7s (the next closest variant), and at 80 AP stimuli, *e*GCaMP^2+^ achieved 1.82x greater response than GCaMP6s (the next closest variant) (**Figure 5B, C; Supp. Table 6B, C**). These results were recapitulated in saturation responses, where the average ΔF/F_0_ response to 40 mM KCl was 1938% for cells expressing *e*GCaMP^2+^ (**Figure 5E; Supp. Table 6E**). This ΔF/F_0_ is 2x greater than those observed in GCaMP6s, the sensor that saw the second-greatest responses to 40 mM KCl (**Figure 5E; Supp. Table 6E**). While the KCl saturation responses were quantified using the cell body, the proximal projections in *e*GCaMP^2+^ maintained >1000% fluorescent increases (**Figure 5F**). At 80 AP trains, both *e*GCaMP and *e*GCaMP^+^ achieved higher fluorescent response amplitudes than the previously published fast variants GCaMP6f and jGCaMP8f (**Figure 5C; Supp. Table 6C**). These results are compounded by both *e*GCaMP and *e*GCaMP^+^ achieving 10 AP half decay times (τ_1/2_) of 1.17s and 0.74s for each variant, respectively. These decay times are faster than jGCaMP8f’s, whose 10 AP half decay time was 1.49s (**Figure 5D; Supp. Table 6D**). Furthermore, *e*GCaMP’s decay kinetics was 8x faster than jGCaMP7s, highlighting the ability of the ensemble to correctly predict the single point mutation’s functional effect (**Figure 5D; Supp. Table 6**).

## Discussion

Incorporating machine learning into our engineering pipeline enabled us to efficiently identify new GCaMP variants with enhanced fluorescent responses and kinetics. We achieved impressive predictive performance in the cross-validation phase by using an ensemble of three regressor models, encoding our dataset with amino acid characteristics, and focusing solely on sequence inputs for learning. These predictive capabilities translated to the *in vitro* space, where many *in silico* predicted characteristics accuratly reflected the mutant’s true performance. As a result of these engineering efforts, we identified three new variants, *e*GCaMP, *e*GCaMP^+^, and *e*GCaMP^2+^.

While the constructs presented here have not been previously described, clues from the literature may explain the impact of these mutations. For example, Residue L317 is known to be involved in extensive hydrophobic interactions between CaM and CBP^27^. Each mutation at L317 that the ensemble proposed is capable of forming hydrogen bonds which may destabilize the CaM and CBP interactions, accelerate kinetics, and altering ΔF/F_0_. Our retrospective analysis revealed previous GCaMP variants that contained a 317E/H/K/N mutation had decreased fluorescent capability compared to jGCaMP7s, which the ensemble learned from the variant library (**Supp. Fig. 4A**). Interestingly, the ensemble identified potential meaningful interactions between residues 52 and 317, both of which are involved in the interaction between CaM and CBP, as all of these variants contained an Alanine at residue 52 (**Supp. Fig. 4B**). Consequently, when we tested the L317H variant in jGCaMP7f, which contains A52, we observed the loss of fluorescence that the model predicted and mirrored previous findings from the Dana *et al.* 2019 study (**Supp. Fig. 4C**). These locations may constitute a promising target for further mutation library studies.

The impressive dynamic range of the Q305D mutation in *e*GCaMP^2+^ may result from intraprotein interactions within CaM. One possible explanation is that the decreased R-group length in the Q305D mutation requires a more substantial conformational change to form the hydrogen bond with residue Y380 (**Suppl. Fig. 3E**). The resulting conformational change may have downstream effects on both the cp-GFP/CaM linker (**Figure 4A**) and on residue R381, which faces inward toward the chromophore (**Suppl. Fig. 3E**). Hence, the dramatic effects of this mutation on the ΔF/F_0_ suggest a collaborative role between the cp-GFP/CaM linker and inward loop of CaM in stabilizing the phenol/phenolate transition of the chromophore^29–31^.

We made several critical design decisions while forming this methodology, such as our encoding method, chosen models, ensemble, and devotion to sequence-only inputs. Dataset encoding is a crucial step in model training as it determines the underlying patterns on which the generalizations are formed^12^. For this reason, we encoded the sequence with biophysical properties underlying the amino acids in each position to form meaningful learning patterns. We derived our AA property datasets from the online repository AAINDEX^32^; however, other similar online databases do exist^33^. Encoding with the property matrices improved the cross-validation R^2^ value by an average of 20% over one-hot encoded or label-encoded libraries (**Supp. Fig 2C**).

Ensembling ML models (i.e., considering the input from multiple models) is advantageous as no singular model is perfectly optimized to perform all tasks^34^. We consider inputs from a random forest regressor (RFR), a K-neighbors regressor (KNR), and a multi-layer perceptron network regressor (MPNR). Decision tree learning methods, such as RFRs, are computationally efficient models well suited for small training libraries, such as the variant library, making them a strong foundation within our ensemble’s learning^12^. KNRs are computationally demanding but simple^35^, where KNR’s similarity metric can capture the degree of variability between the performances of nearly identical sequences. The similarity metric highlights residues whose mutation led to large differences in the targeted biophysical property. MPNRs are deep-learning models capable of extracting high-level features from the data, making them useful for identifying key residues or properties that lead to the observed biophysical response^12^. The three selected models have diverse learning strategies and make different assumptions about the data, which is important when ensembling. When the predictions from each model are ensembled, the cross-validation predictive accuracy matches or improves the sole contributor’s performance (**Supp. Fig 2C**).

While structural insights guided the engineering of previously published GCaMPs, we developed the ensemble pipeline to be structure agnostic. This design consideration was crucial, as we aim to engineer subsequent GEFIs using this pipeline without relying on molecular structures. Due to the exclusion of structure information, extrapolation outside of the observed sequence space may be difficult. This tool is best suited for data generalization and exploration within a sequence space with only minor variations from the training dataset, such as point mutations at tested residues. However, one could incorporate spatial information from crystal structures or structure prediction tools in the ensemble’s learning to aid extrapolation in the future.

The machine learning ensemble used in this study has demonstrated an impressive capacity to guide fluorescent biosensor engineering. The ensemble’s predictions helped identify variants with high dynamic ranges and fast decay kinetics while highlighting clusters of impactful residues for each biophysical property, which may be further exploited by mutation library-based high-throughput screening. These findings illustrate the ensemble’s ability to guide engineering efforts and improve experimental efficiency. Moreover, since our model’s learning is based solely on the sequence-function relationship and all contributor model optimization is unbiased, the final ensemble platform can be broadly applied to any genotype-to-phenotype mutation library. Applying this ML platform to mutation studies of proteins with quantifiable output characteristics, including other protein sensors, has the potential to accelerate the engineering of these proteins.

## Acknowledgments

S.J.W. was supported by DGE-2140004. A.B. was supported by The Brain Research Foundation, UW Royalty Research Fund, UW ISCRM IPA, NIGMS R01 GM139850-01, P30 DA048736-01-Pilot. The research received additional support from the UW NAPE Center and ISCRM Shared Equipment.

## Material requests

Plasmids for eGCaMP+ and eGCaMP2+ can be obtained directly from Addgene for mammalian expression or subcloning encoded in pCAG backbones (#201147, #201148) and virus production for cre dependent expression encoded in pAAV-ef1a-DIO backbones (#201149, #201150).

## Methods

### Data Preprocessing

The Chen and Dana studies provide a functional characterization of >1000 GCaMP variants that span the GCaMP6 and jGCaMP7 iterations^4, 5^. The experimental conditions from each study were standardized across experiments, allowing a direct comparison of the GCaMP mutation’s properties^36^. Each study normalized the results to base constructs for data such as the fluorescent response (ΔF/F_0_, *Eq.1*) to stimuli of 1 AP, 3 AP, 10 AP, 160 AP, and decay half-time after 10 AP. To cross-compare mutation libraries, we re-normalized the *Chen et al.* 2013 dataset such that GCaMP6s was 1.0 for all metrics. The authors linked the functional ability of each variant to a primary key identifier and the identities of the mutations within each variant. The list of mutations was relative to either GCaMP3 or GCaMP6s for *Chen et al. 2013* and *Dana et al. 2019,* respectively. To generate a dataset compatible with ML algorithms, we replaced the list of mutations with a Pandas DataFrame containing one column per residue. The resultant data structure comprised 453 columns: one column containing the primary key identifiers present in the parent datasets, 451 columns corresponding to the sequence of GCaMP, and the final column containing each variant’s empirically derived performance. The mutations that occurred in each variant were reflected in their respective sequence positions within the DataFrame. Any duplicated variants that were present were isolated, and their responses were averaged before compiling them back into the variant library. This duplicate data consideration ensures that each variant only occurs once in the final variant library and ameliorates instances of data leakage between train and test data.

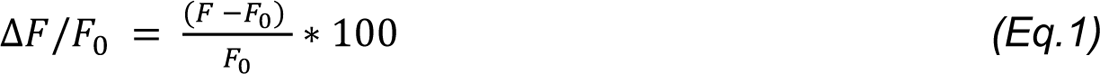

The resultant dataset is the basis for our dependent and independent variables used to train our ML algorithms. Within the variant library used for model training, the independent variable consists of the sequence of each mutation. The dependent variable is the fluorescent response (1AP ΔF/F_0_) or kinetics capability (τ_1/2_). However, because the sequence is a series of string-type values, the complexities of the identities of each amino cannot be understood by the algorithms. The sequences need to be encoded with quantitative values. Dataset encoding can be performed in several capacities: label encoding, one-hot encoding, or by adding functional information. Within our label encoding, we randomly assigned an integer value to each amino acid and replaced each residue label in the GCaMP sequence with the dummy label. For one-hot encoding, the full extent of possible residues at each position is considered in a boolean manner (20 amino acids x 450 residue positions). The start codon, methionine, is considered a one in column 1-M, where every other 1-x contains a zero. Finally, to perform encoding with functional data, we developed a dictionary of amino acid properties by web scraping the AAINDEX database^32, 37^. AAINDEX consists of matrices that each describe a different AA property (e.g., Size (Dawson, 1972), Polarity (Grantham, 1974), Hydrophobicity (Jones, 1975)). We used the 554 complete property datasets to formulate an unbiased model training paradigm in two steps. The AAINDEX contains 566 datasets of published work’s float type values for various amino acid properties, though only 554 contain a value for all 20 amino acids. These 554 datasets became the amino acid property dictionary we used to encode our variant library. The final variant library used in model training consisted of the fully encoded GCaMP sequence and the variants empirically derived performance capability.

### Generation of the novel variant library

To generate a library of unknown sequences, we performed a single-point saturation of the jGCaMP7s sequence at 75 residue locations. These 75 residues correspond to the 75 residues that contain mutagenesis information in the variant library. The outcome was a novel point-saturation-mutation library that contained 1500 sequences. To ensure each variant was a previously untested sequence, we removed variants that had sequences redundant to any that occurred in the variant library, including any redundancies with jGCaMP7s, such that the final point-saturation-mutation library contained 1423 variants. Specifically, 75 variants were redundant with the base jGCaMP7s sequence, and two variants (jGCaMP7s L317A and jGCaMP7s H78K) were redundant with previously characterized variants. The fluorescence and kinetics ensembles generated predictions of the functional capabilities of the 1423 novel variants in the novel library with jGCaMP7s included as a control. These final predictions serve as the basis for mutations considered for *in vitro* testing.

### Ensemble Training

The learning capabilities of any model are limited when tasked to predict outcomes where the factors underlying response have innumerable contributing factors. Under this assumption, we trained and optimized three regressors that would each contribute to the mutation predictions we tested *in vitro*. Our ultimate goal was to ensemble these weak learners and focus our downstream efforts on mutually agreed upon mutations. The models that we developed were from the pip installable package Scikit Learn in Python 3.8.5 to develop a Random Forest Regressor (RFR), K-Neighbors Regressor (KNR), and a Multi-layer Perceptron Network Regressor (MPNR) (**Supp. Table 7**). The models were trained on the encoded sequence of each variant linked to their empirically derived performance capability. The performance capabilities correspond to their fluorescent response to one AP or half decay time after 10 AP. The data was split into train/test sets at a ratio of 80:20 with a random seed of 42 for downstream optimization efforts. Due to the inherent complexity of the 451-residue feature space of the GCaMP sequence, we performed the ‘Sel ectKBest’ feature selection function found in Scikit Learn to rank the importance of each input feature before model training. This feature selection was critical to reduce the dimensionality of the data and, ultimately, decreasing the required runtime. Optimization of the model was done by grid-search hyperparameter tuning. We used the coefficient of determination (R^2^) and mean squared error (MSE) to track the fitting of each model. Additionally, we optimized each model using the key considerations that govern model performance, such as the number of neighbors in KNR and the number of estimators in RFR. Conditions that lead to the highest R^2^ of the test set were compared between each AA property dataset used for encoding to individually optimize and associate predictive capabilities with the underlying amino acid property. This process was repeated over each of the 554 datasets for the three models (∼1662x). Each model’s top five performing property datasets were advanced to generate predictions on the novel variant library. Each contributor model (5 AA property x 3 Regressor models) forms predictions independently, and the final predictions are the average response from each contributor model for each target attribute (fluorescent response 1AP ΔF/F_0_ or kinetics capability τ_1/2_). The predicted values returned by the ensembles are numeric values originating from a normalized library, making the predictions unitless. For example, smaller numeric values in the fluorescence library would correspond to a predicted decreased fluorescent response, and smaller numeric values in the kinetics library would correspond to a predicted faster decay speed (**see Figure 2**).

### PCA Clustering

Each feature within the data was first scaled using Sklearn’s StandardScaler. We passed the scaled data into Sklearn’s PCA function with no defined number of components. We chose the optimum number of components by finding where the explained variance of the PCA of the data passed 0.8. We reinitialized the PCA with the determined number of principal components and fit the function with the standardized data. We then used the principal component space coordinates to find the ideal number of clusters for K-Means clustering. We determined the ideal number of clusters by using the ‘elbow method’ on the Within Cluster Sum of Square. After finding the clusters, we labeled each input to their K-means-defined cluster.

### Molecular Cloning

Predicted mutations were reflected into the CMV-jGCaMP7s backbone (Addgene ID: 104463) using point-mutation primers ordered from Integrated DNA Technologies (IDT) and PCR amplification with either Q5-polymerase (catalog: M0492L) or Superfi-II polymerase (catalog: 12368010). Amplification of the DNA fragment was verified with agarose gel electrophoresis. Blunt-end DNA circularization was achieved with Kinase, Ligase, and DpnI enzyme (KLD) treatment (New England Biolabs: E0554S). Circularized DNA was transformed into competent *E.Coli* cells (DH5ɑ or TOP10) and grown on agar plates that contain either ampicillin or kanamycin selection antibiotic (50 µg/mL). Upon colony formation, single colonies were picked and grown in 5mL cultures containing LB Broth (Fisher BioReagents; BP9723-2) and selection antibiotic (ampicillin/kanamycin; 50 µg/mL) overnight (37°C, 230 RPM). DNA was isolated using Machery Nagel DNA prep kits (Machery Nagel; 740490.250). Sanger sequencing (Genewiz; Seattle, WA) of the isolated plasmid DNA was used to confirm the presence of the intended mutation.

Genes encoding the GCaMP variants were cloned into a CAG-driven backbone, pCAG-Archon1-KGC-EGFP-ER2-WPRE (Addgene; #108423), using Gibson assembly (New England Biolabs; E2621L). All subsequences were verified with Sanger sequencing (Genewiz; Seattle, Wa).

### Acetylcholine Assays

HEK293 cells were cultured in Dulbecco’s Modified Eagle Medium + GlutaMAX (Gibco; 10569-010) supplemented with 10% fetal bovine serum (Biowest; S1620). When cultures reached 85% confluency, the cultures were seeded at 100,000 cells/well or 50,000 cells per well in 24-well and 48-well plates, respectively. 24 hours after cell seeding, the cells were transfected using Lipofectamine3000 (Invitrogen; L3000015) at 1000 ng of DNA per well of a 24-well plate, according to the manufacturer’s instructions. 48 hours post-transfection, the plates were prepared for imaging by washing and then replacing culturing media volume with imaging solution (Tyrode’s pH = 7.33; 125mM NaCl, 2mM KCl, 2 mM CaCl_2_, 2 mM MgCl_2_, 30 mM Dextrose, 25 mM HEPES (triple supplemented with 1% Glutamax (Gibco; 35050-1), 1% Sodium Pyruvate (GIBCO; 11360-070), and 1% MEM Non-Essential Amino Acids (Gibco; 11140-050)). Crystalline power Acetylcholine Chloride (Alfa Aesar; L02168.14) was resuspended into imaging solution (Tyrode’s pH = 7.33; 125mM NaCl, 2mM KCl, 2 mM CaCl_2_, 2 mM MgCl_2_, 30 mM Dextrose, 25 mM HEPES) into 2x the desired final concentration. During imaging, 1:1 volumes of the acetylcholine-tyrodes imaging solution were hand-pipetted into the bath volume to bring the final acetylcholine concentration to the desired concentration. Imaging was performed on a sCMOS camera (Photometrics Prime95B) on an epifluorescent microscope (Leica DMI8) using a 20X objective (Leica HCX PL FLUOTAR L 20x/0.40 NA CORR). A Lumencor Light Engine LED and Semrock Filters (Excitation: FF01-474-27; Emission: FF01-620/35) were used for fluorescence imaging.

### Analysis of Fluorescent Assays

Analysis of Human Embryonic Kidney (HEK293; ATCC Ref: CRL-1573) cell fluorescence imaging data was done by FluorAREA, a custom cloud-based semi-automated time series fluorescence data analysis platform written in Python. First, the cell segmentation quality of the selected Cellpose^38^ model was manually verified. For the segmentation of cells expressing cytosolic fluorescent indicators, model ‘cyto’ was selected as our base model. If the selected Cellpose model was low-performing, we further trained the Cellpose model using the Cellpose 2.0 human-in-the-loop system^39^. Using an “optimized” segmentation model, fluorescence time-series data is extracted for each region of interest. This allows for unbiased extraction of change in cellular fluorescence information for a complete set of experimental samples. Using the raw fluorescence data, % fluorescence change from the baseline (ΔF/F_0_) over time was calculated using *Eq.1*. The signal-to-noise ratio (SNR) was calculated using *Eq.2*.

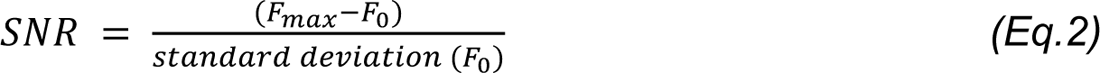

The exponential decay constant (λ) was calculated using Eq.3, where F(t) is the change in fluorescence at a time (t) after the max fluorescence (F_0_) was achieved. Importantly, F_0_ was normalized to 1.0, such that F(t) depicts the change in fluorescence over time, t.

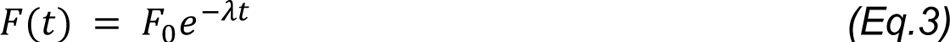

The exponential time constant (τ) was isolated by using the known reciprocal relationship of λ and τ (Eq.4).

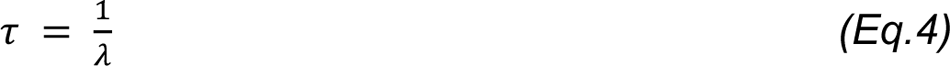

The dynamic range (DR) was defined as the ratio of the max fluorescent intensity to the baseline fluorescent intensity (Eq.5). All ΔF/F_0_, SNR, τ, and DR values were quantified using a custom python script.

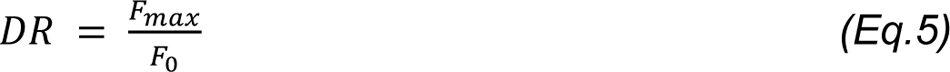

### Isolation of Cortical Neurons

Primary cortical neurons were prepared as previously described^40, 41^. Briefly, 24-well tissue culture plates were coated with matrigel (mixed 1:20 in cold-PBS, Corning; 356231) solution and incubated at 4°C overnight prior to use. Sterile dissection tools were used to isolate cortical brain tissue from P0 rat pups. Tissue was minced until 1mm pieces remained, then lysed in equilibrated (37°C, 5% CO_2_) enzyme (20 U/mL Papain (Worthington Biochemical Corp; LK003176) in 5mL of EBSS (Sigma; E3024)) solution for 30 minutes at 37°C, 5% CO_2_ humidified incubator. Lysed cells were centrifuged at 200xg for 5 minutes at room temperature, and the supernatant was removed before cells were resuspended in 3 mLs of EBSS (Sigma; E3024). Cells were triturated 24x with a pulled Pasteur pipette in EBSS until homogenous. EBSS was added until the sample volume reached 10 mLs prior to spinning at 0.7 rcf for 5 minutes at room temperature. Supernatant was removed, and enzymatic dissociation was stopped by resuspending cells in 5 mLs EBSS (Sigma; E3024) + final concentration of 10 mM HEPES Buffer (Fisher; BP299-100) + trypsin inhibitor soybean (1 mg/ml in EBSS at a final concentration of 0.2%; Sigma, T9253) + 60 µl of fetal bovine serum (Biowest; S1620) + 30 µl 100 U/mL DNase1 (Sigma;11284932001). Cells were washed 2x by spinning at 0.7 rcf for 5 minutes at room temperature and removing supernatant + resuspending in 10 mLs of Neuronal Basal Media (Invitrogen; 10888022) supplemented with B27 (Invitrogen; 17504044) and glutamine (Invitrogen; 35050061) (NBA++). After final wash spin and supernatant removal, cells were resuspended in 10 mLs of NBA++ prior to counting. Just before neurons were plated, matrigel was aspirated from the wells. Neurons were plated on the prepared culture plates at desired seeding density. Twenty-four hours after plating, 1µM AraC (Sigma; C6645) was added to the NBA++ growth media to prevent the growth of glial cells.Plates were incubated at 37°C and 5% CO_2_ and maintained by exchanging half of the media volume for each well with fresh, warmed Neuronal Basal Media (Invitrogen; 10888022) supplemented with B27 (Invitrogen; 17504044) and glutamine (Invitrogen; 35050061) every three days.

### Electrical Field Stimulation

Isolated primary cortical neurons were transfected using the calcium phosphate transfection kit from Sigma Aldrich (Sigma-Aldrich; CAPHOS-1KT). Half of the neuron media was changed 24 hours before transfection, saving the removed conditioned media to add to the neurons after transfection. Reagents were mixed in a ratio of 3 µl CaCl_2_: 24.5 µl H_2_O: 1000 ng DNA before being added dropwise to bubbled 2x HEPES Buffered Saline (30 µl). The final solution was vortexed for 4 seconds and left undisturbed for 20 minutes. The solution was added dropwise to each well of neurons in a 24-well plate and shaken to distribute equally. Neurons were left to incubate for 1 hr at 37^0^C with 5% CO_2_. The cells were rinsed twice with HBSS before adding the conditioned media removed from the day prior and mixed with half-fresh media.

On the day of imaging, ∼24-36 hours post-transfection, cells were washed once with imaging solution and then transferred to E-Stim Tyrode’s (pH = 7.33; 150 mM NaCl, 4 mM KCl, 3 mM CaCl_2_, 1 mM MgCl_2_, 10 mM Dextrose, 10 mM HEPES)^28^. A custom wire holding piece was designed to fit into 48-well plates with silver wires 10 mm apart. 100 mA pulses, with a 3 ms pulse width, were administered at a 10 Hz frequency using a pulse generator (Warner Instruments; SIU-102B), triggered with Sutter Instruments Integrated Patch Amplifier with Patch Panel, time-locked using Igor Pro 8. Imaging was performed with a digital camera (Hamamatsu ORCA-Flash4.0; C11440) at 100ms exposure attached to an epifluorescent microscope (Leica DM IL). The light was generated using a SOLA Light Engine (Lumencor; SOLA SE 5-LCR-SB) with a 488 nm wavelength filter lens. Bulk fluorescence traces were acquired using FIJI imaging software with background subtraction (rolling = 50 stack) and hand-drawn ROIs. The baseline was defined as the first 50 measurements before the event trigger. Max ΔF/F_0_ and decay values were obtained using a custom Python script. Final traces were plotted in Prism9.

### Potassium Chloride Assays

Isolated primary cortical neurons were transfected using the calcium phosphate transfection kit from Sigma Aldrich (Sigma-Aldrich; CAPHOS-1KT). Half of the neuron media was changed 24 hours before transfection, saving the removed conditioned media to add to the neurons after transfection. Reagents were mixed in a ratio of 3 µl CaCl_2_: 24.5 µl H_2_O: 1000 ng DNA before being added dropwise to bubbled 2x HEPES Buffered Saline (30 µl). The final solution was vortexed for 4 seconds and left undisturbed for 20 minutes. The solution was added dropwise to each well of neurons in a 24-well plate and shaken to distribute equally. Neurons were left to incubate for 1 hr at 37^0^C with 5% CO_2_. The cells were rinsed twice with HBSS before adding the conditioned media removed from the day prior and mixed with half-fresh media.

On the day of imaging, ∼24-36 hours post-transfection, cells were washed once with imaging solution, then replaced with imaging solution (Tyrode’s pH = 7.33; 125mM NaCl, 2mM KCl, 2 mM CaCl_2_, 2 mM MgCl_2_, 30 mM Dextrose, 25 mM HEPES (triple supplemented with 1% Glutamax (Gibco; 35050-1), 1% Sodium Pyruvate (Gibco; 11360-070), and 1% MEM Non-Essential Amino Acids (Gibco; 11140-050)). Powdered Potassium Chloride (Sigma; P9541-500G) was diluted in ddH_2_O to a concentration of 2M. This solution was then diluted to 80mM in imaging solution (Tyrode’s pH = 7.33; 125mM NaCl, 2mM KCl, 2 mM CaCl_2_, 2 mM MgCl_2_, 30 mM Dextrose, 25 mM HEPES). During imaging, 1:1 volumes of KCl solution were hand-pipetted into the bath to bring the final KCl concentration to the desired concentration. Imaging was performed on a sCMOS camera (Photometrics Prime95B) on an epifluorescent microscope (Leica DMI8) using a 20X objective (Leica HCX PL FLUOTAR L 20x/0.40 NA CORR). A Lumencor Light Engine LED, and Semrock Filters (Excitation: FF01-474-27; Emission: FF01-620/35) were used for fluorescence imaging. Bulk fluorescence traces were acquired using FIJI imaging software with background subtraction (rolling = 50 stack) and hand-drawn ROIS. The baseline was defined as the first 30 measurements before KCl addition. Max ΔF/F_0_ values were obtained using a custom Python script. Final traces were plotted in Prism9.

