## Supplemental Figures and tables for "Machine Learning Ensemble Directed Engineering of Genetically Encoded Fluorescent Calcium Indicators"

***Supplementary Figure 1: Mutation Scope of Chen & Dana dataset & Train/Test Breakdown***

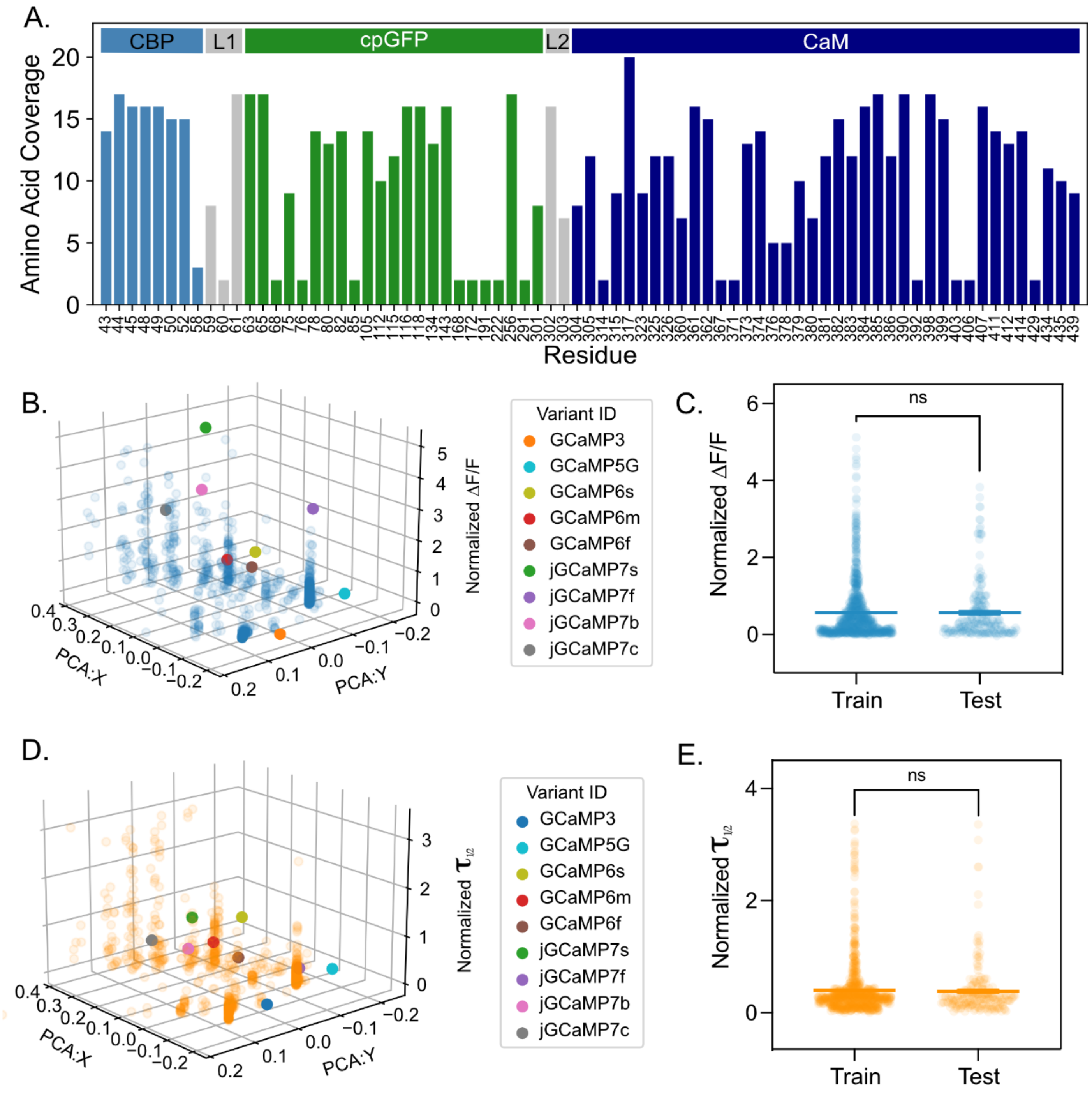


Supplementary Figure 1: *Mutation Scope of Chen & Dana dataset & Train/Test Breakdown*

**A.** Bar plot depicts the number of tested amino acids for each residue in the full variant library. Color coding indicates the location within the GCaMP protein, light blue means the residue is in the CBP, gray means the residue is in one of the two linkers, green means the residue is in the cpGFP, and dark blue means the residue is in the CaM (x-axis denotes residue number, bar height indicates # of amino acids).

**B.** 2D principal component analysis (PCA) of sequences contained in the full variant library with the third dimension displaying the normalized $\Delta F/F_0$ of each variant. Published variants are included as differentially colored dots, indicated in the legend.

**C.** Span of Normalized $\Delta F/F_0$ values in the train set (n=862) and the test set (n=216). Each dot indicates one variant, where the line designates the mean and error bars SEM. Average values and distribution did not differ between the two sets. (ns = P-value>0.05, unpaired t-test)

**D.** 2D PCA of sequences contained in the full variant library with the third dimension displaying the normalized $\tau_{1/2}$ of each variant. Published variants are included as differentially colored dots, indicated in the legend.

**E.** Span of Normalized $\tau_{1/2}$ values in the train set (n=862) and the test set (n=216). Each dot indicates one variant, where the line designates the mean and error bars SEM. Average values and distribution did not differ between the two sets. (ns = P-value>0.05, unpaired t-test)

***Supplementary Figure 2: Fluorescence and Kinetics Ensembles Display Amino Acid Propert Preference***

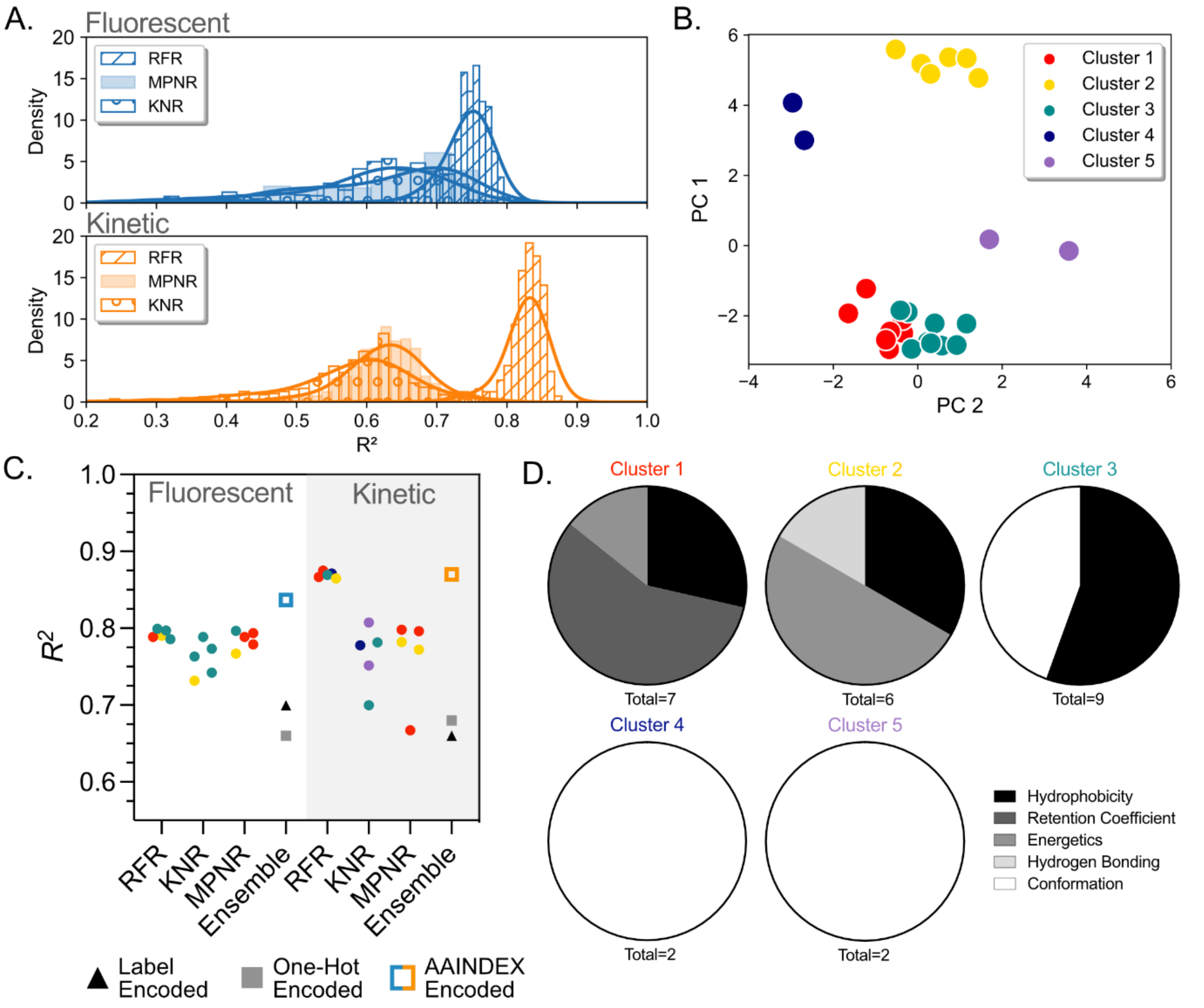


Supplementary Figure 2: *Fluorescence and Kinetics Ensembles Display Amino Acid Property Preference*

- **A.** Kernel density estimates depict the range of the max $R^2$ values saved for each of the 554 amino acid property datasets optimized for both ensembles. Bar patterning designates regressor type. The top five performing property datasets were advanced for downstream analysis.
- **B.** Principal component analysis of the values in the top datasets from each ensemble (2x15 datasets). (number of components = 4, number of clusters = 5).
- **C.** Scatter plot of $R^2$ values from the top five performing amino acid matrices for each regressor type within each ensemble. Color mapping is indicative of PCA cluster identity. Ensemble $R^2$ indicates the final $R^2$ value of predictions from each contributor model after averaging for the indicated encoding method. Models belonging to the fluorescence library are plotted on a transparent background, and models belonging to the kinetic library are plotted on an opaque background.

**D.** Pie-charts depict the amino acid properties that were found in each PCA cluster. Total indicates the number of property matrices within the cluster. Name and name color coordinate with **B./C.**

### *Supplementary Figure 3: In Silico Predictions Indicate Key Residues and Interactions Within GCaMP Protein*

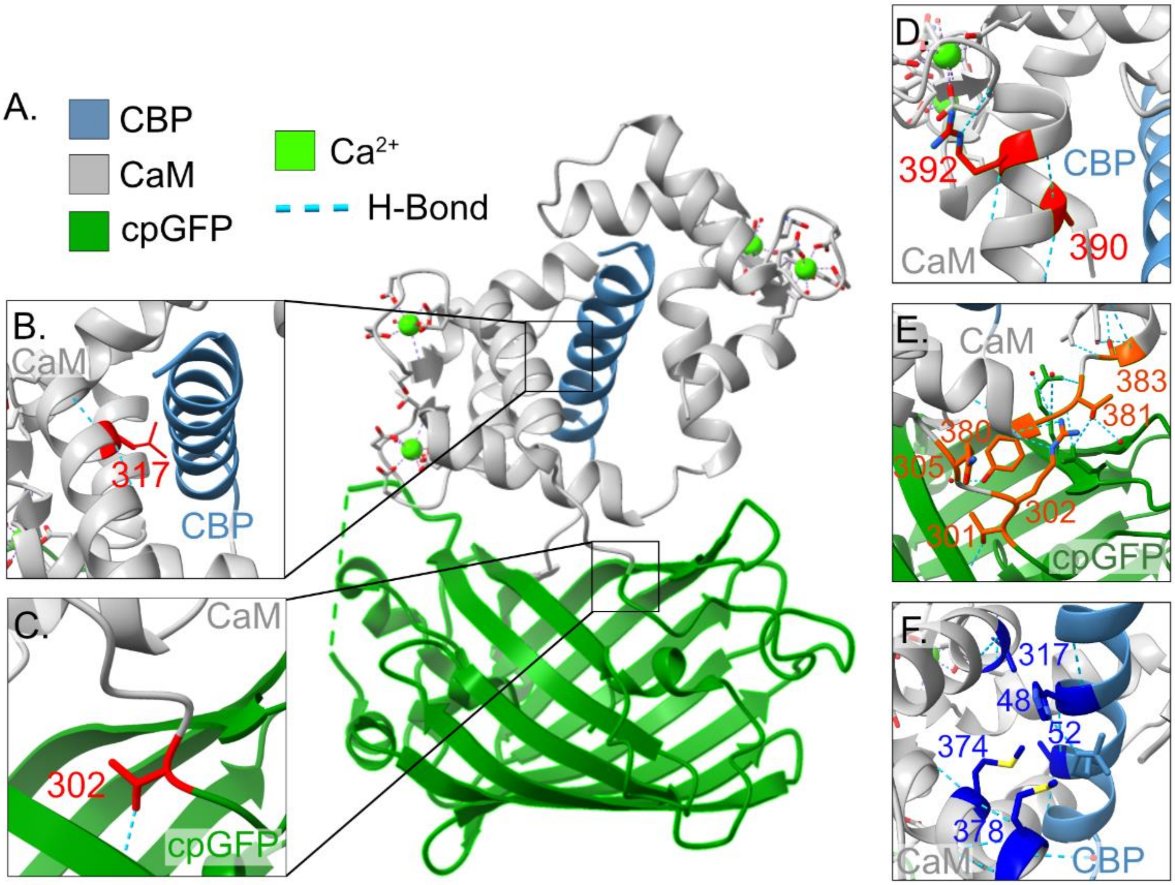


Supplementary Figure 3: *In Silico Predictions Indicate Key Residues and Interactions Within GCaMP Protein*

- **A.** Crystal structure of GCaMP3 D380Y (RCSB: 3SG3), with color mapped CaM (gray), CBP (light blue), cpGFP (dark green), $Ca^{2+}$ (lime green), and hydrogen bonds (light blue dashed lines).
- **B.** Residue A317L rotamer (red) on the interface of CaM (gray) and CBP (light blue).
- **C.** Residue L302 (red), on linker between CaM (gray) and cpGFP (dark green), with hydrogen bonds (light blue dashed lines).
- **D.** Residue A390 (red, left), interfacing with the EF-hand motif, and G392 (red, right) interfacing with CBP (light blue), with color-mapped CaM (gray).
- **E.** Representative image of residues Y380, R381, T383, L302, P303, and Q305 (dark orange) proximity on the crystal structure with color-mapped CaM (gray) and cpGFP (dark green).
- **F.** Representative image of residues K48, V52, L317, M374, and M378 (dark blue) proximity on the crystal structure with color-mapped CaM (gray) and CBP (light blue).

### *Supplementary Figure 4: Retroactive Assessment of Ensemble Predictions Compared to In Vitro Behaviors*

A.

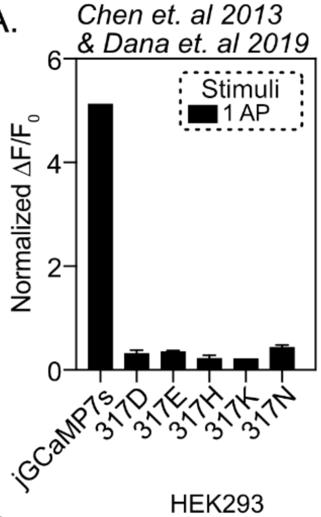


B.

| Index | 52 AA | 317 AA | 374 AA | $\Delta F/F_0$: 1 AP |
|---|---|---|---|---|
| 10.1473 | V | L | M | 5.12 |
| 10.1035 | A | H | M | 0.15 |
| 10.1259 | A | H | M | 0.28 |
| 67 | A | E | M | 0.09 |
| 361 | A | E | M | 0.21 |
| 362 | A | E | M | 0.14 |
| 500 | A | E | M | 0.29 |
| 608 | A | E | M | 0.13 |
| 609 | A | E | M | 0.15 |
| 610 | A | E | M | 0.26 |
| 611 | A | E | M | 0.15 |
| 612 | A | E | M | 0.29 |
| 613 | A | E | M | 0.74 |
| 630 | A | E | M | 0.33 |
| 631 | A | E | M | 0.24 |
| 632 | A | E | M | 0.21 |
| 633 | A | E | M | 0.11 |
| 642 | A | E | M | 0.42 |
| 644 | A | E | M | 0.28 |
| 645 | A | E | M | 0.22 |
| 646 | A | E | M | 0.32 |
| 647 | A | E | M | 0.31 |
| 648 | A | E | M | 0.21 |
| 649 | A | E | M | 0.43 |
| 663 | A | E | M | 0.12 |
| 664 | A | E | M | 0.24 |
| 684 | A | E | M | 0.16 |
| 685 | A | E | M | 0.20 |
| 686 | A | E | M | 0.17 |
| 687 | A | E | M | 0.15 |
| 688 | A | E | M | 0.13 |
| 689 | A | E | M | 0.10 |
| 690 | A | E | M | 0.16 |
| 691 | A | E | M | 0.15 |
| 692 | A | E | M | 0.07 |
| GCaMP6f | A | E | M | 0.67 |
| 10.916 | A | E | M | 0.27 |

| Index | 52 AA | 317 AA | 374 AA | $\Delta F/F_0$: 1 AP |
|---|---|---|---|---|
| 10.1083 | A | E | M | 0.61 |
| 10.1087 | A | E | M | 0.74 |
| 10.1088 | A | E | M | 0.65 |
| 10.1097 | A | E | M | 0.27 |
| 10.1098 | A | E | M | 0.64 |
| 10.1112 | A | E | M | 0.28 |
| 10.1113 | A | E | M | 0.76 |
| 10.1117 | A | E | M | 0.43 |
| 10.1118 | A | E | M | 0.43 |
| 10.1119 | A | E | M | 0.42 |
| 10.1124 | A | E | M | 0.49 |
| 10.1125 | A | E | M | 0.29 |
| 10.1126 | A | E | M | 0.34 |
| 10.1127 | A | E | M | 0.9 |
| 10.1128 | A | E | M | 0.32 |
| 10.1136 | A | E | M | 0.6 |
| 10.1182 | A | E | M | 0.66 |
| 10.1186 | A | E | M | 0.31 |
| 10.1206 | A | E | M | 0.26 |
| 10.1207 | A | E | M | 0.28 |
| 10.1212 | A | E | M | 0.16 |
| 10.1217 | A | E | M | 0.59 |
| 10.1227 | A | E | M | 0.74 |
| 10.1228 | A | E | M | 0.21 |
| 10.1229 | A | E | M | 0.51 |
| 10.123 | A | E | M | 0.78 |
| 10.1236 | A | E | M | 0.64 |
| 10.124 | A | E | M | 0.14 |
| 10.1243 | A | E | M | 0.34 |
| 10.1251 | A | E | M | 0.5 |
| 10.1253 | A | E | M | 0.32 |
| 10.1257 | A | E | M | 0.01 |
| 10.1258 | A | E | M | 0.59 |
| 10.923 | A | N | M | 0.38 |
| 10.1166 | A | N | M | 0.48 |
| 10.92 | A | K | M | 0.21 |

C.

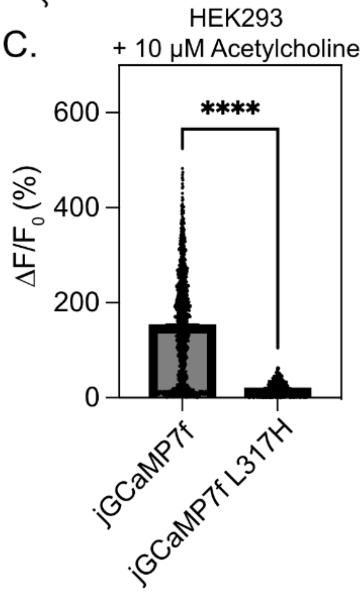


Supplementary Figure 4: *Retroactive Assessment of Ensemble Predictions Compared to In Vitro Behaviors*

- **A.** Normalized fluorescent responses to indicated stimuli in cultured neurons, obtained from the Chen *et al. 2013* and Dana *et al. 2019* variant library (bars depict mean + SEM (if applicable).
- **B.** Table of values contained in A. Values are derived from the Chen *et al. 2013* and Dana *et al. 2019* variant library, rows contain the identification number of the cataloged variant (Index), the variants' amino acid identities at residues 52, 317, and 374 (52 AA, 317 AA, 374 AA), and the normalized $\Delta F/F_0$ at 1 AP ($\Delta F/F_0$: 1AP). jGCaMP7s is included in the first row as it's Dana *et al.* 2019 derived identity (10.1473).
- **C.** Max fluorescent responses obtained from listed variants expressed in HEK293 cells and stimulated with acetylcholine. (n = number of cells quantified; bars depict mean + SEM, **** = <0.0001 (unpaired t-test)). [jGCaMP7f = 150.3 ± 3.5 (n=1036); jGCaMP7f L317H = 17.05 ± 0.65 (n=498)].

**_Supplementary Figure 5: Combinatorial Mutation Biophysical Characteristics and Basis for Mutation Transfer_**

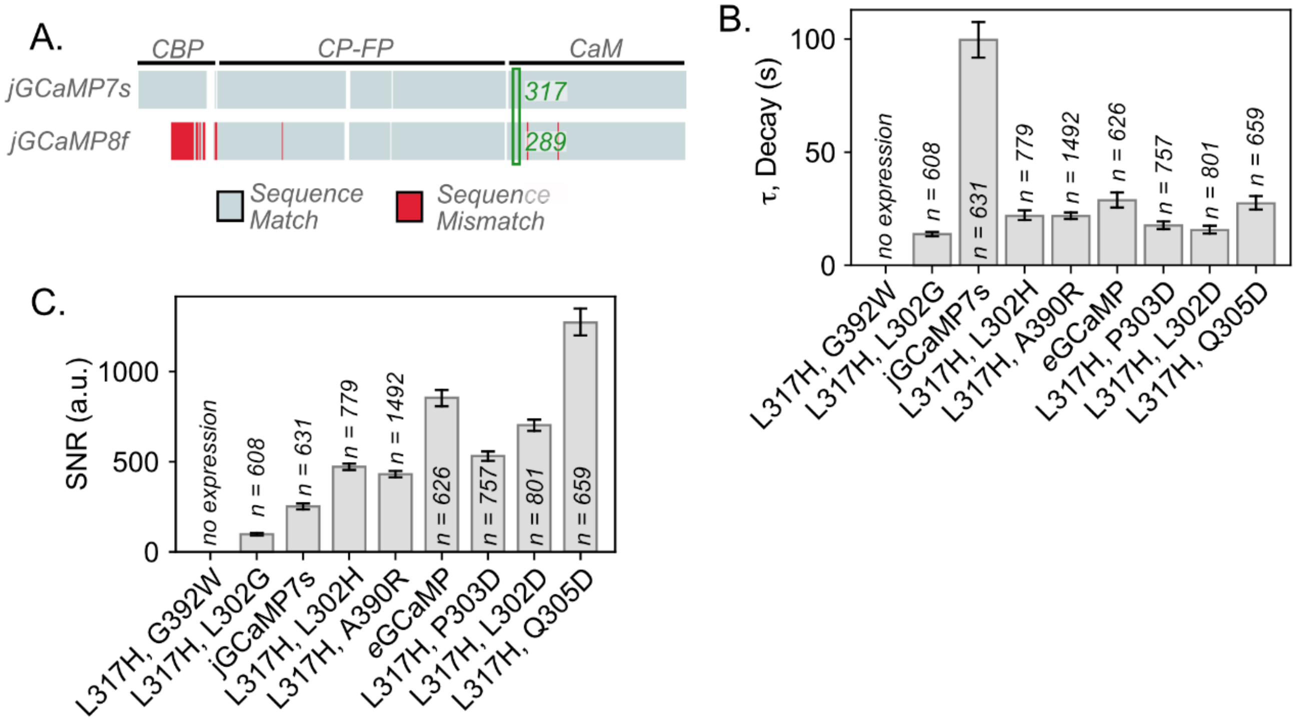


Supplementary Figure 5: *Combinatorial Mutation Biophysical Characteristics and Basis for Mutation Transfer*

- **A.** Sequence alignment of jGCaMP7s and jGCaMP8f. Light gray indicates identical sequence alignment, and red indicates sequence dissimilarities. Breaks indicate a missing sequence portion caused by additional sequence portions in other constructs. The physical location of the residue L317 in jGCaMP7s is designated with the green box and the residue number of the matching location is included as green text over the sequence. Text along the top of the sequence depicts the physical location in the GCaMP protein: CBP, CaM, or circularly permuted fluorescent protein (cpFP).
- **B.** Decay values (**$\tau$**, tau, *Eq.4*) obtained from each combinatorial mutant of jGCaMP7s expressed in HEK293 cells and stimulated with 10 µM acetylcholine. Mutations are sorted according to $\Delta F/F_0$ performance (n = the number of cells quantified; bars depict mean + bootstrapped 95% ci[42]).
- **C.** Signal-to-noise ratio (SNR, *Eq.4*) obtained from each combinatorial mutant of jGCaMP7s expressed in HEK293 cells and stimulated with 10 µM acetylcholine. Mutations are sorted according to $\Delta F/F_0$ performance (n = the number of cells quantified; bars depict mean + bootstrapped 95% ci[42]).

#### ***Supplementary Figure 6: Ratiometric Comparison of eGCaMP Variants' Performance Against Published Variants***

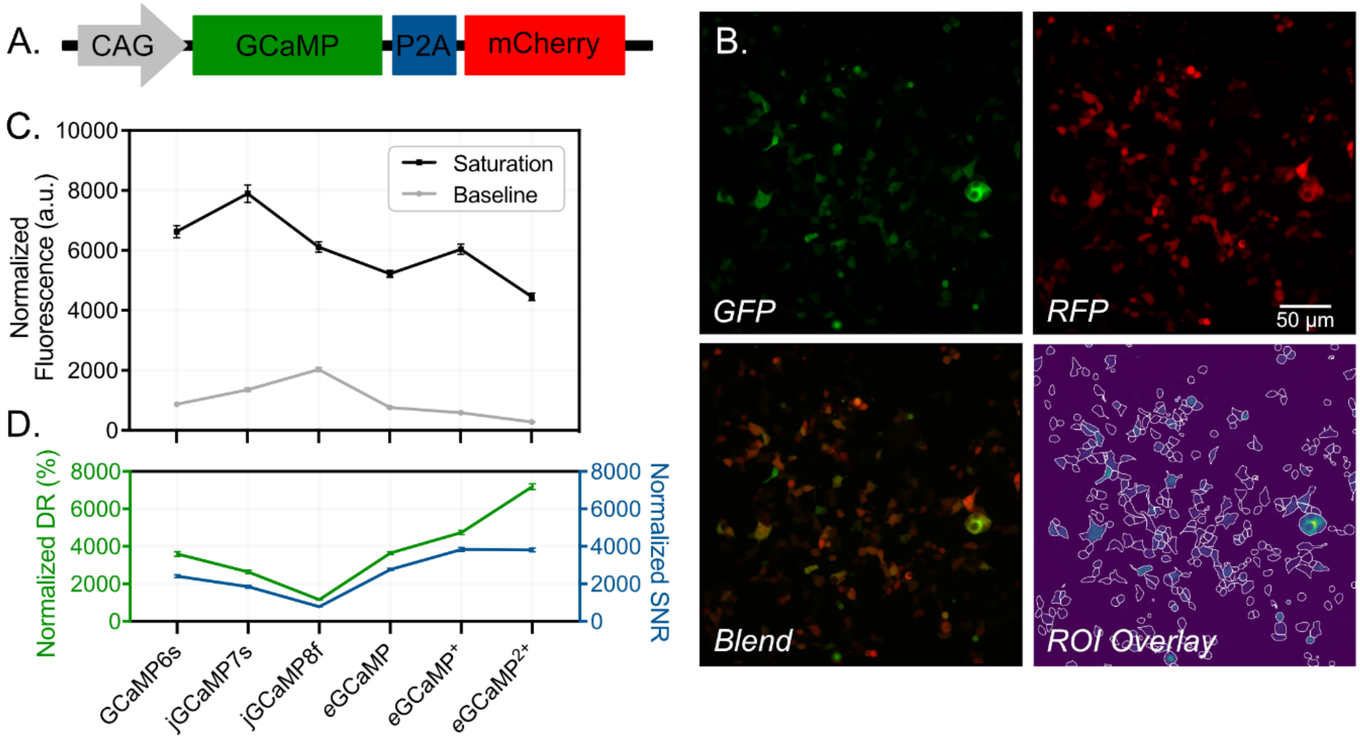


Supplementary Figure 6: *Ratiometric analysis of baseline fluorescence for eGCaMP, eGCaMP$^{+}$, and eGCaMP$^{2+}$*

- **A.** GCaMP variants GCaMP6s, jGCaMP7s, jGCaMP8f, *e*GCaMP, *e*GCaMP$^{+}$, *e*GCaMP$^{2+}$ were transformed into a CAG driven vector, with a mCherry control fluorophore added after a self-cleaving motif (P2A) within the reading frame.
- **B.** Representative images taken with 488 nm wavelength excitation (GFP), 585nm wavelength excitation (RFP), an overlap of the two channels depicting the ratio of GFP intensity to RFP intensity (Blend), and the Cellpose derived ROIs used for analysis (ROI overlay). The scale bar depicts 50µm.
- **C.** Normalized fluorescence intensity of each ratiometric variant (x-axis shared with D.) at baseline (gray) and saturation conditions (black). Scatter points depict mean value and error bars depict SEM.
- **D.** Normalized dynamic range (DR, green, left y-axis) and signal-to-noise ratio (SNR, blue, right y-axis) for each ratiometric variant (x-axis). Scatter points depict mean value and error bars depict SEM.

### Supplemental Tables

#### *Supplementary Table 1: Fluorescence Library Encoding Dataset Information*

| ***Fluorescence*** | | | | |
|---|---|---|---|---|
| Model | AAINDEX | $R^2$ | Descriptor | Cluster # |
| RFR | ROSM880102 | 0.80 | Side chain hydropathy, corrected for solvation (Roseman, 1988) | 2 |
| | MANP780101 | 0.797 | Average surrounding hydrophobicity (Manavalan-Ponnuswamy, 1978) | 3 |
| | KANM800104 | 0.796 | Average relative probability of inner beta-sheet (Kanehisa-Tsong, 1980) | 3 |
| | JURD980101 | 0.795 | Modified Kyte-Doolittle hydrophobicity scale (Juretic et al., 1998) | 3 |
| | MEEJ810102 | 0.795 | Retention coefficient in NaH2PO4 (Meek-Rossetti, 1981) | 1 |
| MPNR | BROC820101 | 0.797 | Retention coefficient in TFA (Browne et al., 1982) | 1 |
| | ZIMJ680105 | 0.787 | RF rank (Zimmerman et al., 1968) | 1 |
| | FAUJ830101 | 0.783 | Hydrophobic parameter pi (Fauchere-Pliska, 1983) | 1 |
| | CIDH920104 | 0.768 | Normalized hydrophobicity scales for alpha/beta-proteins (Cid et al., 1992) | 3 |
| | BULH740101 | 0.766 | Transfer free energy to surface (Bull-Breese, 1974) | 2 |
| KNR | KANM800104 | 0.78 | Average relative probability of inner beta-sheet (Kanehisa-Tsong, 1980) | 3 |
| | LIFS790102 | 0.77 | Conformational preference for parallel beta-strands (Lifson-Sander, 1979) | 3 |
| | MANP780101 | 0.77 | Average surrounding hydrophobicity (Manavalan-Ponnuswamy, 1978) | 3 |
| | BASU050101 | 0.76 | Interactivity scale obtained from the contact matrix (Bastolla et al., 2005) | 3 |
| | MIYS990101 | 0.75 | Relative partition energies derived by the Bethe approximation. (Miyazawa-Jernigan, 1999) | 2 |

Supplementary Table 1: *Fluorescence library encoding dataset information.*

The table contains the top five performing AAINDEX datasets for each model (RFR, MPNR, KNR). The ‘AAINDEX’ column pertains to the property dataset's unique identifier within the AAINDEX repository. ‘$R^2$’ is the performance score of the model when trained on the 80% train set and validated with the 20% test set. ‘Descriptor’ is the description of the property obtained from AAINDEX. ‘Cluster #’ is derived from the principle component analyses on all property datasets, with additional information in Supplementary Figure 2.

### *Supplementary Table 2: Kinetics Library Encoding Dataset Information*

| ***Kinetics*** | | | | |
|---|---|---|---|---|
| Model | AAINDEX | $R^2$ | Descriptor | Cluster # |
| RFR | ZASB82010 | 0.88 | Dependence of partition coefficient on ionic strength (Zaslav sky et al., 1982) | 1 |
| | QIAN880130 | 0.88 | Weights for coil at the window position of -3 (Qian-Sejnowski , 1988) | 4 |
| | RADA880101 | 0.88 | Transfer free energy from chx to wat (Radzicka-Wolfenden, 1988) | 1 |
| | NADH010103 | 0.87 | Hydropathy scale based on self-information values in the tw o-state model (16% accessibility) (Naderi-Manesh et al., 200 1) | 3 |
| | GUYH850104 | 0.87 | Apparent partition energies calculated from Janin index (Gu y, 1985) | 2 |
| MPNR | BULH740101 | 0.79 | Transfer free energy to surface (Bull-Breese, 1974) | 2 |
| | FAUJ880110 | 0.79 | Number of full nonbonding orbitals (Fauchere et al., 1988) | 2 |
| | ZIMJ680105 | 0.78 | RF rank (Zimmerman et al., 1968) | 1 |
| | MEEJ800101 | 0.78 | Retention coefficient in HPLC, pH7.4 (Meek, 1980) | 1 |
| | ROSM880101 | 0.77 | Side chain hydropathy, uncorrected for solvation (Roseman, 1988) | 2 |
| KNR | PALJ810103 | 0.77 | Normalized frequency of beta-sheet from LG (Palau et al., 1 981) | 3 |
| | PALJ810107 | 0.77 | Normalized frequency of alpha-helix in all-alpha class (Palau et al., 1981) | 5 |
| | PALJ810112 | 0.77 | Normalized frequency of beta-sheet in alpha/beta class (Pal au et al., 1981) | 3 |
| | LEVM780103 | 0.77 | Normalized frequency of reverse turn, with weights (Levitt, 1 978) | 4 |
| | QIAN880105 | 0.76 | Weights for alpha-helix at the window position of -2 (Qian-S ejnowski, 1988) | 5 |

Supplementary Table 2: *Kinetics library encoding dataset information.*

The table contains the top five performing AAINDEX datasets for each model (RFR, MPNR, KNR). The 'AAINDEX' column pertains to the property dataset's unique identifier within the AAINDEX repository. '$R^{2}$' is the performance score of the model when trained on the 80% train set and validated with the 20% test set. 'Descriptor' is the description of the property obtained from AAINDEX. 'Cluster #' is derived from the principle component analyses on all property datasets, with additional information in Supplementary Figure 2.

### Supplementary Table 3: Descriptive Statistics of Ensemble Prediction Screen Results

A.

| ΔF/F (%) | | | |
|---|---|---|---|
| Variant | Mean | 95% CI | ROIs Measured (n) |
| L317N | 1224.53 | [1188.62, 1257.82] | 608 |
| L317E | 1239.16 | [1195.84, 1282.91] | 562 |
| L317K | 436.49 | [397.03, 477.46] | 145 |
| L317H | 932.7 | [897.68, 967.47] | 626 |
| L302D | 519.48 | [492.15, 549.41] | 541 |
| L302R | 258.63 | [248.38, 271.61] | 769 |
| L302G | 322.53 | [309.5, 335.76] | 707 |
| L302H | 255.27 | [246.09, 264.53] | 851 |
| L302C | 276.18 | [265.04, 287.55] | 707 |
| A390Y | 149.82 | [142.29, 157.56] | 606 |
| A390R | 906.15 | [884.21, 927.9] | 882 |
| P303D | 620.06 | [598.72, 641.78] | 787 |
| jGCaMP7s | 489.93 | [469.53, 510.23] | 631 |
| P303W | 716.15 | [691.97, 740.69] | 635 |
| P303F | 889.13 | [863.36, 914.67] | 1072 |
| G392F | 932.41 | [896.26, 969.28] | 450 |
| G392W | 1119.84 | [1085.01, 1154.64] | 573 |

B.

| Tau (s) | | | |
|---|---|---|---|
| Variant | Mean | 95% CI | ROIs Measured (n) |
| A390Y | 124.72 | [116.65, 133.5] | 606 |
| L302C | 181.63 | [170.57, 193.2] | 707 |
| L302D | 139.24 | [127.55, 151.71] | 541 |
| L302R | 100.29 | [93.8, 107.2] | 769 |
| A390R | 66.75 | [64.23, 69.38] | 882 |
| L302G | 218.24 | [204.23, 232.75] | 707 |
| L302H | 132.81 | [127.34, 138.61] | 851 |
| P303F | 118.52 | [112.54, 125.0] | 1072 |
| P303W | 94.61 | [89.19, 100.68] | 635 |
| jGCaMP7s | 99.6 | [92.02, 107.83] | 631 |
| G392F | 70.51 | [65.82, 75.64] | 450 |
| G392W | 65.17 | [60.42, 70.23] | 573 |
| L317H | 28.75 | [25.54, 32.31] | 626 |
| P303D | 113.24 | [107.3, 119.51] | 787 |
| L317K | 4.65 | [4.26, 5.08] | 145 |
| L317N | 28.94 | [25.49, 33.0] | 608 |
| L317E | 35.16 | [31.7, 39.14] | 562 |

C.

| SNR (a.u.) | | | |
|---|---|---|---|
| Variant | Mean | 95% CI | ROIs Measured (n) |
| jGCaMP7s | 251.74 | [234.83, 269.59] | 631 |
| L302C | 138.67 | [131.24, 146.11] | 707 |
| L302D | 319.8 | [297.64, 342.55] | 541 |
| L302G | 428.67 | [405.3, 452.19] | 707 |
| L302H | 146.09 | [136.89, 155.66] | 851 |
| L302R | 98.1 | [92.87, 104.14] | 769 |
| P303D | 455.22 | [429.31, 482.85] | 787 |
| P303F | 357.71 | [341.91, 373.36] | 1072 |
| P303W | 343.9 | [325.42, 363.0] | 635 |
| L317E | 810.13 | [753.73, 866.7] | 562 |
| L317K | 133.79 | [117.73, 151.03] | 145 |
| L317H | 854.35 | [808.94, 899.55] | 626 |
| L317N | 586.23 | [541.01, 636.35] | 608 |
| A390R | 559.12 | [533.85, 585.68] | 882 |
| A390Y | 41.99 | [39.03, 45.06] | 606 |
| G392F | 471.71 | [441.75, 502.39] | 450 |
| G392W | 590.15 | [557.76, 623.86] | 573 |

D.

| Performance Score (a.u.) | | | |
|---|---|---|---|
| Variant | Mean | 95% CI | ROIs Measured (n) |
| jGCaMP7s | 3.83 | [3.51, 4.15] | 631 |
| L302C | 1.22 | [1.11, 1.34] | 707 |
| L302D | 4.02 | [3.6, 4.47] | 541 |
| L302G | 3.65 | [3.09, 4.45] | 707 |
| L302H | 1.46 | [1.34, 1.59] | 851 |
| L302R | 1.68 | [1.43, 2.06] | 769 |
| P303D | 5.43 | [5.03, 5.85] | 787 |
| P303F | 4.42 | [4.17, 4.69] | 1072 |
| P303W | 5.02 | [4.67, 5.38] | 635 |
| L317E | 35.48 | [32.71, 38.38] | 562 |
| L317K | 32.71 | [28.56, 37.26] | 145 |
| L317H | 54.49 | [51.17, 57.92] | 626 |
| L317N | 34.89 | [32.0, 37.85] | 608 |
| A390R | 10.4 | [9.84, 10.98] | 882 |
| A390Y | 0.51 | [0.46, 0.56] | 606 |
| G392F | 9.31 | [8.5, 10.13] | 450 |
| G392W | 12.84 | [11.83, 13.94] | 573 |

Supplementary Table 3: *Descriptive Statistics of Ensemble Prediction Screen Results*

Tables containing the information displayed in Figures 3**B,C,D,E**. Tables contain the construct (Variant), the mean response (**A.** ΔF/F (%), **B.** Tau (s), **C.** SNR (a.u), **D.** SNR/Tau (a.u.)), the 95% confidence interval (95% CI), and the number of samples (ROIs Measured (n)).

### *Supplementary Table 4: Descriptive Statistics of Combinatorial Mutation Screen Results*

A.

| ΔF/F (%) | | | | |
|---|---|---|---|---|
| Variant | Acetylcholine Stimuli (µM) | Mean | 95% CI | ROIs Measured (n) |
| L317H, L302G | 10 | 401.8 | [387.48, 416.08] | 608 |
| jGCaMP7s | 10 | 489.93 | [469.11, 510.71] | 631 |
| L317H, L302H | 10 | 758.06 | [738.7, 777.22] | 779 |
| L317H, A390R | 10 | 892.11 | [875.99, 908.27] | 1492 |
| *e* GCaMP (L317H) | 10 | 932.7 | [897.84, 967.47] | 626 |
| L317H, P303D | 10 | 942.58 | [916.89, 968.52] | 757 |
| L317H, L302D | 10 | 1098.63 | [1070.71, 1126.54] | 801 |
| L317H, Q305D | 10 | 2407 | [2335.38, 2478.47] | 659 |

B.

| Tau (s) | | | | |
|---|---|---|---|---|
| Variant | Acetylcholine Stimuli (µM) | Mean | 95% CI | ROIs Measured (n) |
| L317H, L302G | 10 | 13.82 | [12.91, 14.78] | 608 |
| jGCaMP7s | 10 | 99.6 | [92.15, 107.49] | 631 |
| L317H, L302H | 10 | 21.86 | [19.91, 24.12] | 779 |
| L317H, A390R | 10 | 21.83 | [20.57, 23.24] | 1492 |
| *e* GCaMP (L317H) | 10 | 28.75 | [25.57, 32.35] | 626 |
| L317H, P303D | 10 | 17.61 | [16.02, 19.49] | 757 |
| L317H, L302D | 10 | 15.59 | [14.03, 17.49] | 801 |
| L317H, Q305D | 10 | 27.39 | [24.62, 30.51] | 659 |

C.

| SNR (a.u.) | | | | |
|---|---|---|---|---|
| Variant | Acetylcholine Stimuli (µM) | Mean | 95% CI | ROIs Measured (n) |
| L317H, L302G | 10 | 97.7 | [91.52, 104.22] | 608 |
| jGCaMP7s | 10 | 251.74 | [234.96, 269.24] | 631 |
| L317H, L302H | 10 | 471.92 | [454.95, 489.27] | 779 |
| L317H, A390R | 10 | 430.55 | [413.14, 448.84] | 1492 |
| *e* GCaMP (L317H) | 10 | 854.35 | [809.76, 901.75] | 626 |
| L317H, P303D | 10 | 531.58 | [504.94, 558.54] | 757 |
| L317H, L302D | 10 | 702.85 | [672.35, 733.8] | 801 |
| L317H, Q305D | 10 | 1271.48 | [1200.07, 1343.99] | 659 |

D.

| SNR/Tau (a.u.) | | | | |
|---|---|---|---|---|
| Variant | Acetylcholine Stimuli (µM) | Mean | 95% CI | ROIs Measured (n) |
| L317H, L302G | 10 | 10.35 | [9.56, 11.17] | 608 |
| jGCaMP7s | 10 | 3.83 | [3.51, 4.15] | 631 |
| L317H, L302H | 10 | 35.66 | [33.73, 37.61] | 779 |
| L317H, A390R | 10 | 27.58 | [26.26, 29.05] | 1492 |
| *e* GCaMP (L317H) | 10 | 54.49 | [51.21, 57.96] | 626 |
| L317H, P303D | 10 | 44.43 | [41.78, 47.16] | 757 |
| L317H, L302D | 10 | 62.97 | [59.92, 66.04] | 801 |
| L317H, Q305D | 10 | 74.62 | [69.97, 79.53] | 659 |

Supplementary Table 4: *Descriptive Statistics of Combinatorial Mutation Screen Results*

Tables containing the information displayed in Figures 4**E,F** and Supplementary Figure 5**B,C**. Tables contain the construct (Variant), the concentration of the acetylcholine stimulus (Acetylcholine Stimuli (µM)), the mean response ((**A.** ΔF/F (%), **B.** Tau (s), **C.** SNR (a.u), **D.** SNR/Tau (a.u.)), the 95% confidence interval (95% CI), and the number of samples (ROIs Measured (n)).

### *Supplementary Table 5: Descriptive Statistics of Acetylcholine Concentration Curve Results*

A.

| Concentration (µM) | Variant | Mean ΔF/$F_0$ (%) | SEM | Samples (n) |
|---|---|---|---|---|
| 10 | GCaMP6s | 542.06 | 16.3 | 1218 |
| 10 | GCaMP6f | 580.06 | 19.75 | 776 |
| 10 | jGCaMP7s | 309.67 | 8.69 | 1401 |
| 10 | jGCaMP7f | 776.22 | 8.83 | 1883 |
| 10 | jGCaMP8s | 189.90 | 4.34 | 1129 |
| 10 | jGCaMP8m | 257.06 | 4.63 | 1514 |
| 10 | jGCaMP8f | 268.50 | 5.21 | 1288 |
| 10 | eGCaMP | 724.49 | 14.36 | 1618 |
| 10 | eGCaMP2+ | 1567.52 | 24.54 | 1924 |
| 10 | eGCaMP+ | 1162.09 | 21.91 | 1467 |

B.

| Concentration (µM) | Variant | Mean ΔF/$F_0$ (%) | SEM | Samples (n) |
|---|---|---|---|---|
| 5 | GCaMP6s | 529.34 | 14.26 | 1275 |
| 5 | GCaMP6f | 582.71 | 14.46 | 1355 |
| 5 | jGCaMP7s | 332.85 | 10.67 | 910 |
| 5 | jGCaMP7f | 743.66 | 7.57 | 1867 |
| 5 | jGCaMP8s | 194.31 | 4.17 | 1143 |
| 5 | jGCaMP8m | 250.46 | 5.08 | 1203 |
| 5 | jGCaMP8f | 256.31 | 4.69 | 1358 |
| 5 | eGCaMP | 715.41 | 14.6 | 1527 |
| 5 | eGCaMP2+ | 1526.99 | 26.47 | 1448 |
| 5 | eGCaMP+ | 1171.96 | 23.31 | 1161 |

C.

| Concentration (µM) | Variant | Mean ΔF/$F_0$ (%) | SEM | Samples (n) |
|---|---|---|---|---|
| 1 | GCaMP6s | 389.18 | 18.02 | 785 |
| 1 | GCaMP6f | 447.79 | 19.66 | 584 |
| 1 | jGCaMP7s | 271.94 | 11.82 | 686 |
| 1 | jGCaMP7f | 542.35 | 9.69 | 1524 |
| 1 | jGCaMP8s | 188.81 | 5.06 | 749 |
| 1 | jGCaMP8m | 203.67 | 6.75 | 760 |
| 1 | jGCaMP8f | 200.35 | 5.39 | 1065 |
| 1 | eGCaMP | 497.50 | 15.62 | 1000 |
| 1 | eGCaMP2+ | 1389.23 | 29.65 | 1125 |
| 1 | eGCaMP+ | 698.07 | 21.61 | 996 |

D.

| Concentration (µM) | Variant | Mean ΔF/$F_0$ (%) | SEM | Samples (n) |
|---|---|---|---|---|
| 0.75 | GCaMP6s | 329.41 | 15.49 | 804 |
| 0.75 | GCaMP6f | 429.50 | 14.13 | 1016 |
| 0.75 | jGCaMP7s | 268.25 | 9.28 | 1006 |
| 0.75 | jGCaMP7f | 477.93 | 9.81 | 1361 |
| 0.75 | jGCaMP8s | 135.62 | 4.28 | 805 |
| 0.75 | jGCaMP8m | 173.83 | 5.27 | 983 |
| 0.75 | jGCaMP8f | 164.20 | 3.92 | 1671 |
| 0.75 | eGCaMP | 474.77 | 11.93 | 1666 |
| 0.75 | eGCaMP2+ | 1079.67 | 26 | 1278 |
| 0.75 | eGCaMP+ | 648.48 | 19.95 | 1034 |

E.

| Concentration (µM) | Variant | Mean ΔF/$F_0$ (%) | SEM | Samples (n) |
|---|---|---|---|---|
| 0.5 | GCaMP6s | 290.84 | 14.57 | 826 |
| 0.5 | GCaMP6f | 293.70 | 18.37 | 657 |
| 0.5 | jGCaMP7s | 223.09 | 11.05 | 846 |
| 0.5 | jGCaMP7f | 442.49 | 9.13 | 1574 |
| 0.5 | jGCaMP8s | 151.18 | 6.43 | 462 |
| 0.5 | jGCaMP8m | 157.39 | 5.01 | 1035 |
| 0.5 | jGCaMP8f | 128.58 | 4.69 | 1100 |
| 0.5 | eGCaMP | 350.12 | 12.21 | 1175 |
| 0.5 | eGCaMP2+ | 1033.06 | 25.58 | 1353 |
| 0.5 | eGCaMP+ | 644.64 | 19.25 | 1121 |

F.

| Concentration (µM) | Variant | Mean ΔF/$F_0$ (%) | SEM | Samples (n) |
|---|---|---|---|---|
| 0.25 | GCaMP6s | 105.54 | 15.68 | 302 |
| 0.25 | GCaMP6f | 128.22 | 12.6 | 491 |
| 0.25 | jGCaMP7s | 156.55 | 11.77 | 453 |
| 0.25 | jGCaMP7f | 214.02 | 7.26 | 1577 |
| 0.25 | jGCaMP8s | 102.76 | 5.92 | 477 |
| 0.25 | jGCaMP8m | 91.13 | 4.29 | 935 |
| 0.25 | jGCaMP8f | 79.28 | 4.02 | 934 |
| 0.25 | eGCaMP | 249.34 | 13.68 | 784 |
| 0.25 | eGCaMP2+ | 546.50 | 18.21 | 1471 |
| 0.25 | eGCaMP+ | 313.19 | 15.23 | 976 |

G.

| Concentration (µM) | Variant | Mean ΔF/$F_0$ (%) | SEM | Samples (n) |
|---|---|---|---|---|
| 0.1 | GCaMP6s | 78.84 | 14.76 | 474 |
| 0.1 | GCaMP6f | 50.79 | 13.63 | 356 |
| 0.1 | jGCaMP7s | 77.28 | 9.16 | 381 |
| 0.1 | jGCaMP7f | 81.91 | 4.79 | 1385 |
| 0.1 | jGCaMP8s | 58.18 | 5.5 | 300 |
| 0.1 | jGCaMP8m | 35.94 | 3.16 | 626 |
| 0.1 | jGCaMP8f | 40.50 | 3.1 | 937 |
| 0.1 | eGCaMP | 71.75 | 6.5 | 896 |
| 0.1 | eGCaMP2+ | 205.69 | 14.34 | 807 |
| 0.1 | eGCaMP+ | 145.32 | 10.61 | 808 |

H.

| Concentration (µM) | Variant | τ, Decay (s) | SEM | Samples (n) |
|---|---|---|---|---|
| 5 | GCaMP6s | 72.31 | 3.06 | 940 |
| 5 | GCaMP6f | 24.58 | 1.03 | 1357 |
| 5 | jGCaMP7s | 100.9 | 4.27 | 676 |
| 5 | jGCaMP7f | 45.97 | 1.42 | 2088 |
| 5 | jGCaMP8s | 30.44 | 0.69 | 1379 |
| 5 | jGCaMP8m | 32.28 | 1.19 | 1325 |
| 5 | jGCaMP8f | 33.52 | 1.13 | 1457 |
| 5 | eGCaMP | 34.02 | 1.49 | 1564 |
| 5 | eGCaMP2+ | 39.02 | 1.74 | 1509 |
| 5 | eGCaMP+ | 18.13 | 0.79 | 1297 |

Supplementary Table 5: *Descriptive Statistics of Acetylcholine Concentration Curve Results*

Tables containing the information displayed in Figures 4**G,H**. (**A.,B.,C.,D.,E.,F.,G.**) Tables contain the concentration of the acetylcholine stimulus (Concentration (µM)), the construct (Variant), the mean response (Mean ΔF/$F_0$ (%)), the standard error of the mean (SEM), and the number of samples (Samples (n)). (**H.**) The table contains the concentration of the acetylcholine stimulus (Concentration (μM)), the construct (Variant), the speed of off-decay (Tau Off (s)), the standard error of the mean (SEM), and the number of samples (Samples (n)).

***Supplementary Table 6: Descriptive Statistics of Primary Neuron Recordings***

A.

| 1 AP | | | | | | | |
|---|---|---|---|---|---|---|---|
| Construct | GCaMP6s | GCaMP6f | jGCaMP7s | jGCaMP8f | *e* GCaMP | *e* $GCaMP^{+}$ | *e* $GCaMP^{2+}$ |
| Number of values | 33 | 48 | 102 | 72 | 56 | 49 | 47 |
| Mean | 3.911 | 2.626 | 6.252 | 8.438 | 2.783 | 3.026 | 10.1 |
| Std. Deviation | 3.197 | 5.021 | 11.51 | 18.88 | 2.845 | 3.702 | 16.25 |
| Std. Error of Mean | 0.5564 | 0.7247 | 1.139 | 2.225 | 0.3802 | 0.5289 | 2.371 |

B.

| 10 AP | | | | | | | |
|---|---|---|---|---|---|---|---|
| Construct | GCaMP6s | GCaMP6f | jGCaMP7s | jGCaMP8f | *e* GCaMP | *e* $GCaMP^{+}$ | *e* $GCaMP^{2+}$ |
| Number of values | 49 | 55 | 134 | 50 | 82 | 77 | 53 |
| Mean | 43.77 | 20.99 | 48.53 | 31.94 | 37.81 | 30.54 | 113.8 |
| Std. Deviation | 53.41 | 34.86 | 57.84 | 35.49 | 50.59 | 64.68 | 207.4 |
| Std. Error of Mean | 7.63 | 4.701 | 4.996 | 5.019 | 5.587 | 7.371 | 28.49 |

C.

| 80 AP | | | | | | | |
|---|---|---|---|---|---|---|---|
| Construct | GCaMP6s | GCaMP6f | jGCaMP7s | jGCaMP8f | *e* GCaMP | *e* $GCaMP^{+}$ | *e* $GCaMP^{2+}$ |
| Number of values | 49 | 63 | 111 | 72 | 88 | 86 | 54 |
| Mean | 276.1 | 114.6 | 142.4 | 84.28 | 275.3 | 190.4 | 502.5 |
| Std. Deviation | 252.1 | 173.6 | 126.7 | 64.44 | 243.7 | 193.1 | 370.1 |
| Std. Error of Mean | 36.01 | 21.87 | 12.03 | 7.594 | 25.97 | 20.82 | 50.37 |

D.

| 10 AP Decay | | | | | | | |
|---|---|---|---|---|---|---|---|
| Construct | GCaMP6s | GCaMP6f | jGCaMP7s | jGCaMP8f | *e* GCaMP | *e* $GCaMP^{+}$ | *e* $GCaMP^{2+}$ |
| Number of values | 16 | 44 | 70 | 47 | 75 | 62 | 43 |
| Mean | 4.37 | 0.9547 | 9.479 | 1.49 | 1.168 | 0.7355 | 2.099 |
| Std. Deviation | 2.583 | 0.4496 | 5.487 | 0.9786 | 0.5577 | 0.6476 | 1.595 |
| Std. Error of Mean | 0.6457 | 0.06778 | 0.6558 | 0.1427 | 0.0644 | 0.08225 | 0.2433 |

E.

| 40 mM KCl | | | | | | | |
|---|---|---|---|---|---|---|---|
| Construct | GCaMP6s | GCaMP6f | jGCaMP7s | jGCaMP8f | *e* GCaMP | *e* $GCaMP^{+}$ | *e* $GCaMP^{2+}$ |
| Number of values | 80 | 42 | 121 | 15 | 120 | 29 | 91 |
| Mean | 941.4 | 373.8 | 550.8 | 232.6 | 873.9 | 833 | 1938 |
| Std. Deviation | 448 | 281.5 | 235.3 | 89.03 | 416.8 | 570.7 | 926.6 |
| Std. Error of Mean | 50.09 | 43.43 | 21.39 | 22.99 | 38.04 | 106 | 97.13 |

Supplementary Table 6: *Descriptive Statistics of Primary Neuron Recording*

Tables containing the information displayed in Figures 5**A,B,C,D**. Tables **A.**, **B.**, and **C.** contain the construct (Construct), number of samples (Number of Values), the mean $\Delta F/F_0$ (%) response (Mean), the standard deviation (Std. Deviation), and the standard error of the mean (SEM) at 1 AP, 10 APs and 80 APs, respectively. Table **D.** contains the construct (Construct), number of samples (Number of Values), the mean half decay time (s) (Mean), the standard deviation (Std. Deviation), and the standard error of the mean (SEM) after 10 AP stimuli. Table **E.** contains the construct (Construct), number of samples (Number of Values), the mean $\Delta F/F_0$ (%) response (Mean), the standard deviation (Std. Deviation), and the standard error of the mean (SEM) after 40 mM KCl Stimulus.

### *Supplementary Table 7: Virtual Environment Information*

| Package | Version |
|---|---|
| Python | 3.8.5 |
| Numpy | 1.22.2 |
| Pandas | 1.2.4 |
| Matplotlib | 3.3.4 |
| Scikit-Learn | 0.24.2 |
| Scipy | 1.5.2 |
| Seaborn | 0.11.1 |
| Re | 2.2.1 |

Supplementary Table 7: *Virtual Environment Information*

Table contains the Python libraries used to perform modeling, data analyses, and data visualization.
